# Marine nematodes exhibit widespread symbiosis, novel chemoautotrophy, and evolutionary conservation of holobiont taxa

**DOI:** 10.64898/2026.08.12.744518

**Authors:** Alejandro De Santiago, Min Khant Han, Sarah B. Hargadon, Mirayana Marcellino Barros, Simone Brito De Jesus, Tiago José Pereira, Holly M. Bik

## Abstract

Microbial symbioses drive the evolutionary and functional diversification of eukaryotic clades, from single-celled protists to large invertebrates. However, our knowledge of host-associated assemblages (the “holobiont”) is limited in microscopic animal phyla with a body size <1mm, due to practical challenges such as low biomass and difficult taxonomy of host species. Marine nematodes represent an ideal case study for rapidly advancing our knowledge of bacterial-animal symbioses, representing a globally abundant invertebrate group with strong links to terrestrial and model organism species within the same phylum. Here, we sequenced the holobionts of 220 marine nematodes and generated 815 metagenome-assembled genomes (MAGs) of host-associated bacteria/archaea. Our data indicates that 20-34% of marine nematodes harbor an obligate intracellular symbiont, often with multiple endosymbionts co-occurring within the same host. Three bacterial phyla (Pseudomonadota Bacteroidota, and Verrucomicrobiota) account for three-quarters of all nematode-associated MAGs, and the majority of these holobiont MAGs represent deeply divergent lineages in the prokaryotic tree of life. The Flavobacteriaceae (a core microbiome taxon in *C. elegans* and other terrestrial nematodes), were consistently recovered across phylogenetically diverse marine nematode lineages, suggesting evolutionary conservation of holobiont taxa across marine and terrestrial environments. We also report a novel chemoautotroph family (*Ca.* Thionematobacter) recovered from nematode hosts in both deep-sea and shallow-water habitats, and report the first confirmed instance of *Cardinium* endosymbionts from marine invertebrates. Finally, ∼65% of nematode-associated MAGs are able to degrade chitin, via hexosaminidase, implying that benthic invertebrate holobionts make significant contributions to global carbon cycling. These results underline the importance of evaluating symbiosis in microscopic marine invertebrates, and accelerating our understanding of animal evolution and ecosystem dynamics in vast benthic habitats.

## Introduction

Microbial symbioses drive the evolution and functional diversification of all eukaryotic lineages (González-Pech et al., 2024; Sunagawa et al., 2020), and understanding animal-microbe partnerships is fundamental for understanding animal biology (McFall-Ngai et al., 2013). Microbial symbionts are widely acknowledged to play key roles in the development, nutrition, and evolutionary fitness of their animal hosts (Douglas, 2017; González-Pech et al., 2024; McFall-Ngai et al., 2013), and broad scientific knowledge of animal-microbe symbioses is increasingly critical for understanding systems ecology. Unfortunately, most research efforts on animal-microbe symbiosis have focused on a small subset of taxa. Our knowledge of protist, sponge, cnidarian, and mollusc symbioses have rapidly advanced over the last few decades (Busch et al., 2022; Hughes & Girguis, 2023; Husnik et al., 2021; McCauley et al., 2023; van Oppen & Blackall, 2019; Voolstra et al., 2024), bolstered by coordinated genomics sequencing efforts, taxon-centric collaborations, and data synthesis efforts (McCauley et al., 2023; Vergara-Florez & Duda, 2026; Webster & Thomas, 2016). In contrast, our knowledge of symbiosis in meiofaunal animal phyla (microscopic metazoans with a body size <1mm, such as nematodes, kinorhynchs, gastrotrichs, and, flatworms (Bik et al., 2012; Martínez et al., 2025)) remains largely unexplored.

Marine sediments cover 70% of the Earth’s surface and constitute one of the planet’s largest reservoirs of global biodiversity (Snelgrove, 1999). Meiofaunal animals are numerically dominant on the seafloor and fundamental to many benthic ecosystem processes, driving nutrient cycling, organic matter remineralization, microbial grazing, and energy transfer through marine food webs (Cerca et al., 2018; Martínez et al., 2025). Phylum Nematoda (roundworms) dominates most benthic meiofauna assemblages, with nematode worms typically comprising 70-90% of metazoan life in marine sediments (Ingels et al., 2014). Despite their global dominance, we have a poor understanding of host-microbe interactions in nematodes and most other benthic meiofaunal groups. Most studies to date have focused on characterizing the microbiomes of microscopic invertebrates using low-resolution 16S rRNA metabarcoding approaches (Bellec et al., 2019; Boscaro et al., 2022; Leasi et al., 2024; Schuelke et al., 2018; Vecchi et al., 2018). Although these studies often recover a predictable set of core microbiome taxa and demonstrate that animal microbiomes are distinct from the surrounding environmental microbial assemblages, it has not been possible to determine whether most core bacterial taxa represent true animal symbionts using 16S rRNA datasets alone. For example, although the largest 16S rRNA study to date reported no evidence of phylosymbiosis in an analysis of ∼1,000 meiofaunal specimens across 21 phyla (Boscaro et al., 2022), targeted genome-scale studies of specific bacterial genera such as *Pseudoalteromonas* have subsequently confirmed phylosymbiosis in multiple animal phyla and FISH approaches further revealed the presence of symbiont biofilms in the esophagus and ovaries of marine nematodes (De Santiago et al., 2026); previously, *Pseudoalteromonas* was frequently reported as a core microbiome taxa in 16S microbiome studies of diverse marine invertebrates (Leasi et al., 2024; Pereira et al., 2023; Schuelke et al., 2018; Y.-T. Zhu et al., 2024) and known to play a key role in the larval development and metamorphosis of larger invertebrates (Ericson et al., 2019; Ohdera et al., 2023; Peng et al., 2020). Genome-scale data – spanning both animal hosts and their microbial associates – is clearly needed to advance our understanding of animal-microbe symbiosis, particularly for small-bodied meiofaunal taxa with difficult taxonomy and high biodiversity.

Nematode worms represent an ideal case study for evaluating broad patterns of animal-microbe symbiosis amongst microscopic invertebrates. First, nematodes represent one of the most speciose animal phyla on earth, with an estimated 1-100 million species spread across almost every marine and terrestrial ecosystem on the planet (Lambshead, 1993; Mokievsky & Azovsky, 2002). Second, nematode worms are renown for being “evolutionary commuters”, with an estimated 30 major habitat transitions across the phylum and at least 15 independent origins of parasitic lifestyles (Blaxter & Koutsovoulos, 2015; Holterman et al., 2019). Nematoda is the only major metazoan phylum that has persistently colonized and diversified in marine, freshwater, and terrestrial environments, with major habitat transitions being the rule rather than the exception (marine-terrestrial, deep-sea to shallow-water, free-living to host-associated lifestyles, etc. (Holterman et al., 2019; Schratzberger et al., 2019)). Microbial partnerships are increasingly thought to have played a role in the evolutionary success of nematodes, with microbial horizontal gene transfer (HGT) events being implicated in enhancing nematode foraging and facilitating lifestyle transitions to parasitism (Lo et al., 2024; Mayer et al., 2011). Third, several nematode lineages represent well-known or emerging systems for symbiosis research, including Stilbonematid nematodes with sulfur-oxidizing *Candidatus* Thiosymbion ecosymbionts (Sogin et al., 2020, 2021), *Astomonema spp.* nematodes with a vestigial gut tract containing *Ca.* Thiosymbion endosymbionts (Musat et al., 2007; Sogin et al., 2021), Oncholaimid nematodes harboring mucus-associated symbiont biofilms (Bellec et al., 2018, 2019; De Santiago et al., 2026), plant-parasitic dagger nematodes (*Xiphinema spp.*) hosting *Xiphinematobacter* endosymbionts presumed to be involved in nutritional mutualisms (Brown et al., 2015), and filarial nematodes (vertebrate parasites such as *Brugia spp.*) associated with intracellular *Wolbachia (Setegn et al., 2024)*. Similar to many invertebrates, phylogenetic patterns indicate multiple independent origins of these nematode symbioses which occur in at least five orders within the phylum (Trejo-Meléndez & Contreras-Garduño, 2025; Zimmermann et al., 2016). Finally, advancing our knowledge of nematode symbioses will have important repercussions for model organism research in *Caenorhabditis elegans* and other genetically tractable nematode systems such as *Pristionchus pacificus* (Trejo-Meléndez & Contreras-Garduño, 2025), with experimental work already demonstrating that bacteria associated with marine nematodes can also enhance the development and lifespan of terrestrial nematodes (Xue et al., 2024). Marine nematodes are poorly studied in contrast to their terrestrial and parasitic counterparts, and given the likely marine origin of nematodes and most other invertebrate phyla (Ahmed et al., 2022; Bik et al., 2010; Y.-C. Lee et al., 2023), benthic seafloor habitats likely represent an underexplored reservoir for diverse animal-symbiont partnerships.

In this study, we used a single-specimen sequencing approach to characterize the bacterial/archaeal holobiont associated with individual marine nematodes collected from shallow-water, deep-sea, and polar habitats worldwide. Nematode holobionts represent a low-complexity metagenome, enabling robust recovery of host-associated prokaryotic metagenome-assembled genomes (MAGs) alongside host genome skims (i.e., including traditional DNA barcoding loci for nematode hosts such as 18S rRNA and mitochondrial COI). Single-specimen metagenomes thus offer a more comprehensive view of animal-microbe symbioses, avoiding primer biases associated with traditional 16S microbiome sequencing (Bartoš et al., 2024; Jones et al., 2025; Schloss, 2021; Strube, 2021; Zaiko et al., 2022), enabling recovery of rare holobiont taxa due to higher sequencing depth (Becker & Pushkareva, 2023), and allowing for broad functional characterization of putative symbiont genomes (Villada et al., 2025).

## Results

### Marine nematode holobionts harbor diverse bacterial and archaeal taxa

We generated single-specimen metagenomes from 220 marine nematodes representing most major marine nematode orders from six geographically distinct environments (**Figure 1A**). Our dataset included marine nematode hosts spanning eight taxonomic orders in Phylum Nematoda: Plectida (n = 3), Araeolaimida (n = 6), Monhysterida (n = 21), Desmoscolecida (n = 15), Desmodorida (n = 55), Chromadorida (n = 21), Enoplida (n = 98) and Triplonchida (n = 1). Our sampling strategy incorporated nematodes from the continental slope in the Beaufort Sea, Alaska (n = 10), intertidal sediments in Bodega Bay Harbor, California (n = 3), deep-sea sediments in the hypoxic Santa Monica Basin in Southern California (n = 12), continental shelf sites around Antarctica (n = 18), temperate muddy and sandy shallow-water habitats around Tybee Island, Georgia (n = 58), and intertidal and subtidal tropical coral sand sediments in the Florida Keys (n = 127; **Figure 1A and Table S1**). Within our nematode dataset, 34 specimens were identified as subfamily Stilbonematinae (Desmodorida), chemosynthetic nematodes with sulfur-oxidizing *Ca.* Thiosymbion ectosymbiont bacteria attached to the cuticle (Zimmermann et al., 2016); all Stilbonematid specimens were recovered from the Florida Keys (**Figure 1BC**). Additionally, 42 specimens were identified as family Oncholaimidae (Enoplida), a group of large-bodied, predatory nematodes commonly found in in both deep-sea and shallow-water habitats, where bacterial symbionts are commonly observed in the nematode mouth, esophagus and various locations along the cuticle (Bellec et al., 2018, 2019; De Santiago et al., 2026) (**Figure 1BC**). Nematode morphological identifications were carried out using light microscopy (specimens identified down to family or genus level), and taxonomic assignments of each worm were further evaluated using host 18S rRNA genes recovered from each metagenome. We carried out a manual BLAST search against public databases using the nematode 18S rRNA sequence recovered from each specimen, and taxonomic assignments were refined or corrected if needed using host molecular data. No 18S rRNA genes were able to be recovered from 35 nematode specimens (15.91%), most likely related to the degradation of high molecular weight DNA from frozen samples and/or the inherent difficulty of assembling the host rRNA genes from short-read meteagenomic datasets (potentially due to high levels of intragenomic variation in some nematode taxa (Bik et al., 2013; Pereira et al., 2020; Pereira & Baldwin, 2016). The 18S rRNA reads recovered from single-specimen nematode metagenomes were also used as a checkpoint to assess potential co-recovery of gut contents, parasites, or other eukaryotic microbiome associates. Non-host 18S rRNA sequences were recovered from 40 nematode specimens (18.18%), and the majority of these co-recovered rRNA reads were assigned to protists and fungi (**Figure S1)**.

**Figure 1.**
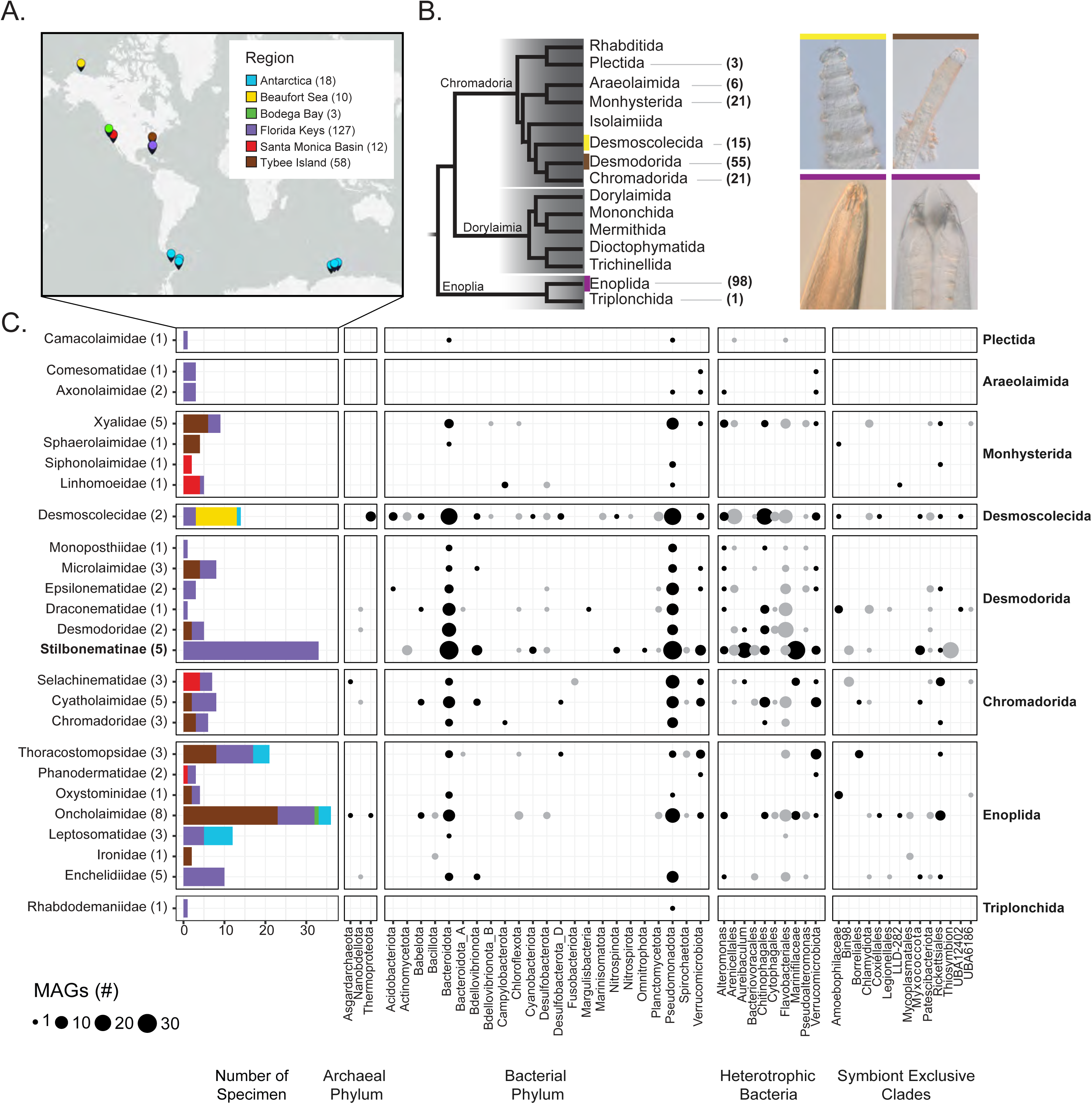
Collection of 220 specimens representing most marine nematode orders. A) Specimens were isolated from six environments ranging from coastal estuarine systems in California to the Antarctic continental shelf. Numbers within parenthesis indicate total specimens recovered from each habitat. B) Cladogram of major nematode lineages, numbers within parenthesis indicate total specimens in each order with micrographs of select representatives from three nematode orders. Dorylaimia is not included in this study as this class is only composed of terrestrial and freshwater nematodes. Bacterial PhyloPics (courtesy of Matt Cook) indicate nematode orders where a symbiotic-exclusive bacterial lineage was recovered (Villada et al., 2025). C) MAGs recovered from each nematode family (left). Each nematode family is grouped by their respective order (right). Chemosynthetic nematodes (members of the subfamily Stilbonematinae that have ectosymbiotic sulfur-oxidizing bacteria *Thiosymbion*) are in bold. Numbers within parenthesis indicate total unique genera. For simplicity, specimens with only family-level classification were counted as one. The bar chart corresponds to the number of specimens in each family whereas the size of each circle corresponds to the number of MAGs recovered.

Individual nematode metagenomes were processed using the MeioBIOME Snakemake pipeline, a new toolkit optimized for the analysis of single-specimen meiofaunal metagenomes which preserves strain-level genomic diversity that may be present in animal holobionts (De Santiago & Bik, 2026). We recovered a total of 815 nematode-associated MAGs, comprising 270 high-quality, 381 medium-quality, and 164 low-quality MAGs from ∼55% of the sampled specimens encompassing 28 bacterial and archaeal phyla (**Figure 1C and Table S2**). We did not recover any MAGs from ∼45% of our nematode specimens. We anticipate that this is likely due to 1) amplification bias of host DNA during REPLI-g whole genome amplification reactions, 2) degraded DNA in nematode samples that were previously frozen or preserved in DESS buffer (Yoder et al., 2006) prior to DNA extraction and amplification, and 3) high-strain level diversity within each specimen, which may prevent accurate binning of prokaryote taxa (in our dataset this was evidenced by a high number of low-quality MAGs from certain nematode specimens, where all binned MAGs consistently contained >10% contamination). Across our entire dataset, three bacterial phyla (*Pseudomonadota*, *Bacteroidota*, and *Verrucomicrobiota*) accounted for 76.81% of all nematode-associated MAGs (**Figure 1C**). The dominance of these three phyla was consistent with the SingleM phyla-level taxonomic report we produced from unassembled metagenomic data (**Supplementary Figure S2**), indicating that the MAGs recovered from our biological samples are a comprehensive assessment of the phylogenetic diversity contained within nematode holobionts. Additionally, 13 archaeal MAGs representing *Asgardarchaeota*, *Nanobdellota*, and *Thermoproteota* were assembled from hosts spanning four nematode orders (Enoplida, Desmodorida, Chromadorida, and Desmoscolecida; **Figure 1C and Table S2**).

Heterotrophic bacteria capable of degrading complex carbohydrates were also common in nematode holobionts, with 30.67% of all the MAGs identified as the orders/families *Chitinophagales*, *Cytophagales*, *Flavobacteriales*, or *Marinifilaceae* (**Figure 1C**). These major heterotrophic bacterial groups were also reliably recovered across phylogenetically distant nematode hosts, suggesting evolutionary conservation of these holobiont taxa (**Figure 1C**). *Pseudoalteromonas* MAGs, which exhibits phylosymbiosis with Oncholaimid nematodes and other diverse marine invertebrate taxa (De Santiago et al., 2026), was sporadically recovered from the dataset, which was consistent with previously reported variable signals outside of family Oncholaimidae. Additionally, 25 *Ca.* Thiosymbion MAGs were recovered from 25 stilbonematid nematodes, with a single *Ca.* Thiosymbion MAG recovered from each host nematode. These *Ca*. Thiosymbion MAGs confirmed that our nematode holobiont dataset was able to recover and correctly classify MAGs corresponding to known bacterial ectosymbionts, and the holobiont patterns of *Ca*. Thiosymbion MAGs (one MAG per Stilbonematid nematode) is in alignment with prior evolutionary studies of these chemosynthetic nematodes (Scharhauser et al., 2020; Zimmermann et al., 2016).

### Nematode holobiont MAGs are undersampled lineages in the prokaryotic tree of life

To further assess the phylogenetic novelty of nematode-associated MAGs, we calculated FastAAI between each MAG and its closest GTDB representative genome (Gerhardt et al., 2025). The median FastAAI value across all prokaryotic MAGs (58.61%) was below the proposed genus-level delimitation threshold of 62.30%, indicating that these nematode-associated genomes are highly divergent from other representative genomes in public databases and likely represent novel genera. (**Figure 2A**). Symbiont-exclusive bacterial clades (defined as bacterial clades where >90% of the lineages are known or computationally predicted obligate intracellular symbionts; (Villada et al., 2025)) recovered from marine nematodes were even more phylogenetically divergent, exhibiting a median FastAAI of 50.53%, which falls below the the proposed family-level delimitation threshold of 53.00% (**Figure 2B)**.

**Figure 2.**
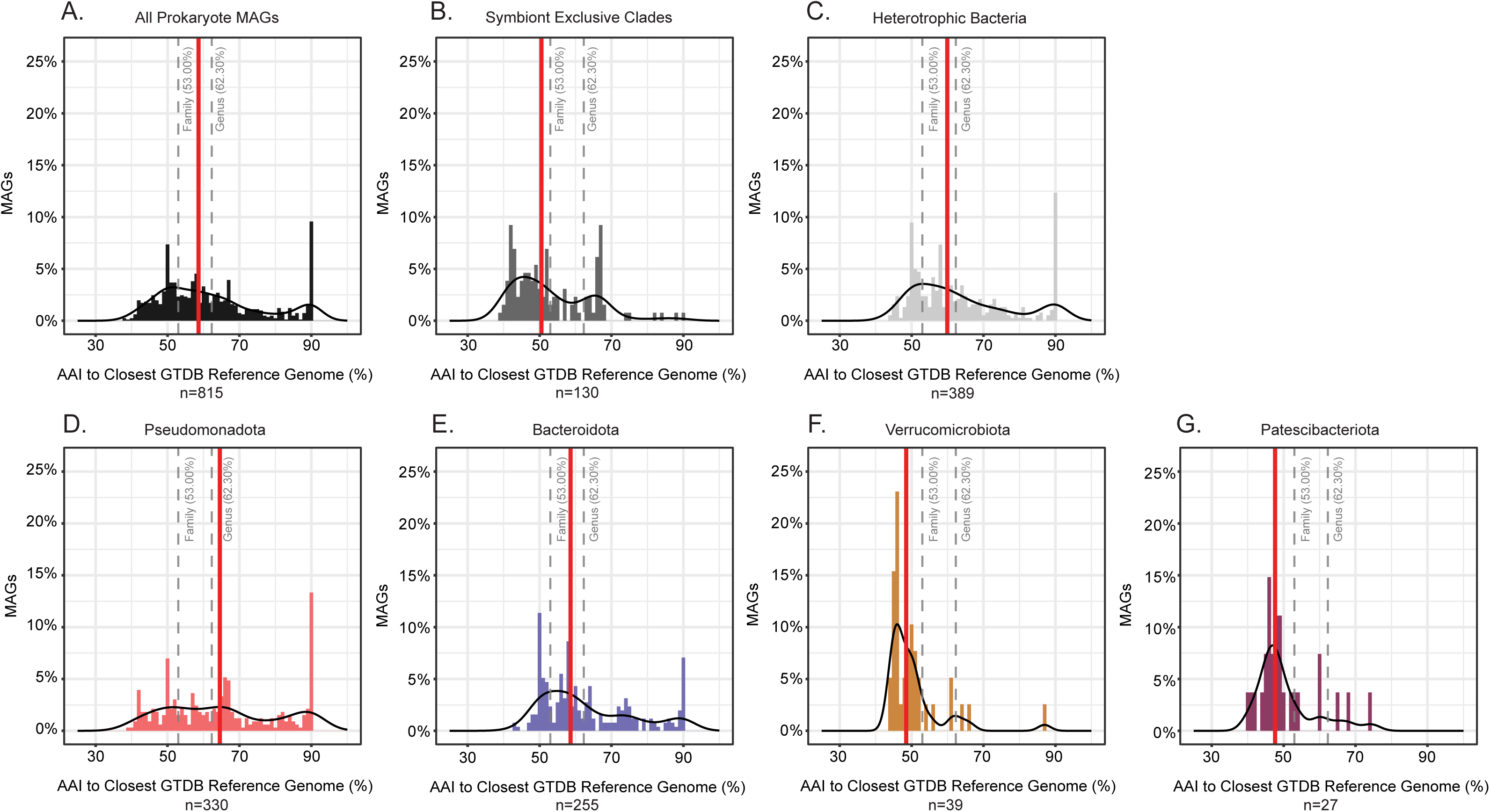
The nematode microbiome is a reservoir of undiscovered microbial lineages. FastAAI (Gerhardt et al., 2025) was compared between bacterial MAGs generated in this study and the closest GTDB reference genome. MAGs were grouped into 3 groups: A) all prokaryote MAGs, B) symbiont-exclusive clades, and C) select heterotrophic bacteria conserved across nematode hosts. FastAAI of the four most common bacterial phylum recovered from nematode holobiont were compared to the closest GTDB reference genome: D) *Pseudomonadota*, E) *Bacteroidota*, F) *Verrucomicrobiota*, and G) *Patescibacteriota.* The grey dash lines indicate the FastAAI typically used to delineate family and genus (Gerhardt et al., 2025). Red lines indicates median FastAAI.

This pattern was also observed across the four most commonly recovered phyla with median FastAAI values of 48.49% for *Pseudomonadota,* 58.53% for *Bacteroidota,* 48.49% for *Verrucomicrobiota, and* 47.62% for *Patescibacteriota*, indicating that >50% of the recovered MAGs are highly divergent from reference genomes and nematode holobiont members likely consist of undescribed deep-branching microbial lineages. These results were consistent across host taxonomy (**Figure S3**), and in line with previous observations that most invertebrate holobiont taxa lack any genus-level genomic representatives in reference databases (Wu et al., 2025). Notably, we recovered 10 of the 15 “most wanted” microbial taxa identified in a recent prokaryotic genome census (**Figure 1C**). The most common “most-wanted” phyla that we recovered in nematode holobionts were *Pseudomonodota* (n=330), *Bacteroidota* (n=257), *Verrucomicrobiota* (n=39), *Patescibacteriota* (n=27), and *Bdellovibrionota* (n=22), representing lineages that are known to contain high levels of phylogenetic diversity yet lack genome representatives for most genera in public databases such as GTDB (Wu et al., 2025).

### Marine nematodes are associated with diverse obligate intracellular symbionts and co-symbiosis is common

To assess the overall prevalence of symbiosis in marine nematodes, we assigned taxonomy to unassembled data using taxonomically informative windows of single-copy genes (SCGs), and subsequently subset the MAGs belonging to symbiont-exclusive clades. Following our prior analysis, symbiont lineages were clades containing ≥90% symbionts, based on a previous study by Villada et al. (2025). Of the 220 nematode holobionts sequenced, 81 had a MAG recovered from a symbiont-exclusive clade (36.82%). Of the nematode samples with symbionts, 31 nematodes (38.27%) had more than one symbiont, and 14 nematodes (17.28%) had more than two symbionts (**Figure S2**). The five most common symbionts were *Patescibacteriota* (n = 44; 20%), *Rickettsiales* (n = 31; 14.09%), *Chlamydiota* (n = 25; 11.36%), *Legionellales* (n = 7; 3.18%), and *Coxiellales* (n = 6; 2.73%). When excluding Patescibacteriota (which are primarily thought to be epibionts of bacteria (Srinivas et al., 2024)), 56 nematode specimens (25.45%) had a symbiont. A total 130 MAGs in our dataset, which were recovered from 54 nematode samples (24.54%), were classified as belonging to a symbiont-exclusive clade. These symbiont MAGs were recovered from 5 of the 8 marine nematode orders that were included in this study (**Figure S1**). However, nematode orders without any symbiont-exclusive MAGs (Plectida, Araeolaimida, and Triplonchida) had low sampling (<6 specimens per order), indicating that increased taxon sampling could further recover MAGs from known bacterial symbiont clades from these undersampled nematode lineages.

Next, we generated phylogenetic trees of a subset of endosymbiont lineages that were well represented in our dataset (also including *Patescibacteriota*) in order to assess their phylogenetic diversity and evolutionary relationships to other known invertebrate symbionts. First, we investigated MAGs belonging to the *Rickettisales,* a bacterial order of ancient obligate intracellular symbionts that have a broad host range (i.e., protists, amoebas, invertebrates species) across diverse environments. For nematode-associated *Rickettisales*, a phylogenetic tree was constructed using GTDB species-level representatives,12 Rickettsiales MAGs, and 11 “Bin98” MAGs recovered from this study (**Figure 3**). Within the *Rickettisales,* we recovered *Anaplasmataceae* MAGs from three nematode hosts (collected from Antarctica and the Florida Keys) which were closely-related to symbionts of chordates and other marine invertebrates (**Figure 3A**), as well as *Midichloriaceae* MAGs from two *Adoncholaimus* (Oncholaimidae) nematodes from Tybee Island, GA and four deep-sea Selachinematidae nematodes from Southern California (**Figure 3B**). The Oncholaimid-associated *Midichloriaceae* formed a sister clade with *Candidatus* Lariskella, arthropod symbionts which have recently been found to infect the ovaries of female Enoplid nematodes (family Thoracostomopsidae) collected from tropical intertidal sands in Okinawa, Japan (Hagenbeek et al., 2026). Furthermore, the *Midichloriaceae* MAGs recovered from deep-sea Selachinematidae nematode hosts were placed as a well-supported monophyletic clade that formed a sister taxon to *Acquirickettsia*, which are known symbionts of other marine invertebrates such as corals (Giannotti et al., 2022). Two other MAGs were classified within the lineages *Rickettsiaceae* and one additional MAG was classified as *UBA1997* (**Figure 3CD**). A single *Rickettsiaceae* MAG was placed in a clade within a monophyletic clade composed of symbionts associated with the ciliate protist genus *Ciliophora*, while the other *Rickettsiaceae* MAG was placed within a clade of aquatic environmental samples. The single UBA1997 MAGs was also placed within a clade containing aquatic environmental samples. Eleven nematode-associated MAGs classified as *Bin98* formed a well-supported phylogenetic clade that included three species-representative genomes classified as WRAU01, a putative family that includes two arthropod symbiont genomes. All Bin98 MAGs in this study were recovered from just two hosts: *Paralaxus sp.* Stilbonematid nematodes from Florida Keys and deep-sea Selachinematidae nematodes from the hypoxic Santa Monica Basin (**Figure 3E**). SingleM taxonomy indicates that *Rickettsiales* is prevalent across all five well-sampled marine nematodes orders in our study, with 7.27% - 26.23% of nematode specimens within each order containing metagenomic signals of *Rickettsiales* (**Figure S1**).

**Figure 3.**
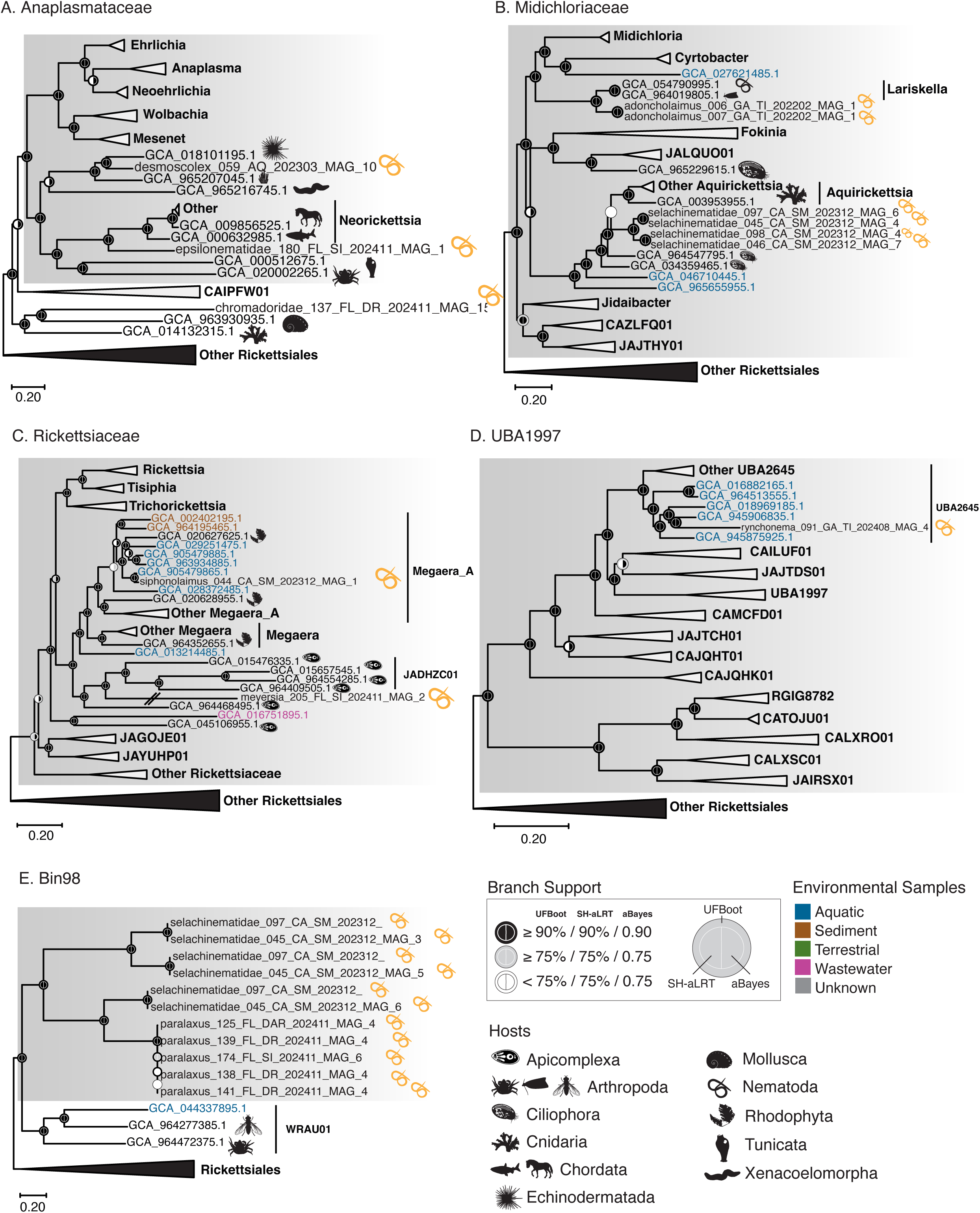
Nematode-associated *Rickettsiales* exhibit broad phylogenetic diversity and expanded host range. A maximum-likelihood phylogenetic tree of *Rickettsiales* was generated with 74 single-copy genes (SCGs) using 640 GTDB species-level representatives, 12 *Rickettsiales* genomes recovered in this study, and 23 *Gammaproteobacteria* genomes to serve as an outgroup were included in the final tree. Focal subtrees of A) *Anaplasmataceae*, B) *Midichloriaceae*, C) *Rickettsiaceae*, D) *UBA1997* were generated by pruning each family from the full phylogenetic tree, with the remaining *Rickettsiales* order shown as a sister reference clade. The *Gammaproteobacteria* outgroup is not shown. E) A maximum-likelihood phylogenetic tree Bin98 was generated with 4541 GTDB alphaproteobacteria family-level representatives, 11 Bin98 genomes recovered in this study, and 23 *Gammaproteobacteria* genomes to serve as an outgroup. Phylopics are used to indicate hosts. Golden nematode phylopics indicate nematode-associate MAGs generated from this study.

Next, we assessed MAGs belonging to the *Chlamydiota,* a diverse clade of predicated obligate endosymbionts that are commonly found in environmental metagenomic sampling of shallow-water and deep-sea sediments (Davison & Hurst, 2023; Lagkouvardos et al., 2014). To assess the diversity of nematode-associated *Chlamydiota* endosymbionts, we constructed a phylogenetic tree using GTDB species-level representatives and seven *Chlamydiota* MAGs constructed from this study (**Figure S4**). Our nematode-associated MAGs were closely related to *Chlamydiota* genomes recovered from hydrothermal vent sediments, wastewater samples, and subsea tunnel biofilms, indicating that MAGs recovered from environmental sequencing efforts may represent undescribed symbionts of microscopic marine invertebrates. Within *Chlamydiota,* we recovered two MAGs classified as JAAKR01 which were placed next to genomes recovered from sediment environmental samples collected near Loki’s Castle hydrothermal vent (**Figure S4**). Two *Chlamydiota* MAGs recovered from Draconematidae (Desmodorida) and *Paralaxus sp.* (subfamily Stilbonematinae) nematodes were classified as JAJFMA01 and SM23-29, respectively (**Figure S4**). Three other MAGs classified as *Simkiniaceae* were recovered from phylogenetically diverse host nematodes spanning three taxonomic orders, including *Daptonema sp.* (Monhysterida), *Meyersia sp.* (Enoplida), and Draconematidae (Desmodorida). The *Chlamydiota* MAGs recovered from *Daptonema* and Draconematidae nematode hosts were closely related to *Simkiniaceae* MAGs recovered from wastewater and biofilms. However, the *Simkiniaceae* MAG recovered from a *Meyersia sp.* nematode was placed into a clade with a sponge symbiont. SingleM taxonomy indicates that *Chlamydiota* is prevalent across all five well-sampled marine nematode orders in our study, and we recovered *Chlamydiota* MAGs from 5.45% - 46.67% of nematodes within each taxonomic order (**Figure S1**). Almost half of marine nematodes in the order Desmoscolecida (7 out of 15 nematode hosts) were associated with at least one *Chlamydiota* MAG.

Next, we assessed nematode-associated MAGs belonging to the *Amoebophilaceae,* a symbiont clade that includes *Cardinium*. Bacteria within the genus *Cardinium* are vertically transmitted endosymbionts that manipulate host reproduction, and have been extensively described from terrestrial arthropods and nematodes; however, to date, no reports of *Cardinium* have been confirmed in any marine invertebrate (Mathieson et al., 2025). To assess the phylogenetic diversity of *Amoebophilaceae*, we constructed a phylogenetic tree containing four *Amoebophilaceae* MAGs recovered in this study alongside GTDB species-level representatives (**Figure 4**). Two MAGs recovered from marine nematodes formed a sister clade with *Cardinium spp.* bacteria associated with terrestrial invertebrates (Mathieson et al., 2025), and this sister clade included *Cardinium* reproductive endosymbionts well described from plant-parasitic terrestrial nematodes (Guo et al., 2026; Tarlachkov et al., 2026). Two additional *Amoebophilaceae* MAGs in our dataset formed a clade with *Amoebophilaceae* from marine and terrestrial environmental samples (**Figure 4**); both of these *Amoebophilaceae* MAGs were recovered from the same nematode, an *Oxystomina sp.* (Enoplida) nematode from Tybee Island, GA.

**Figure 4.**
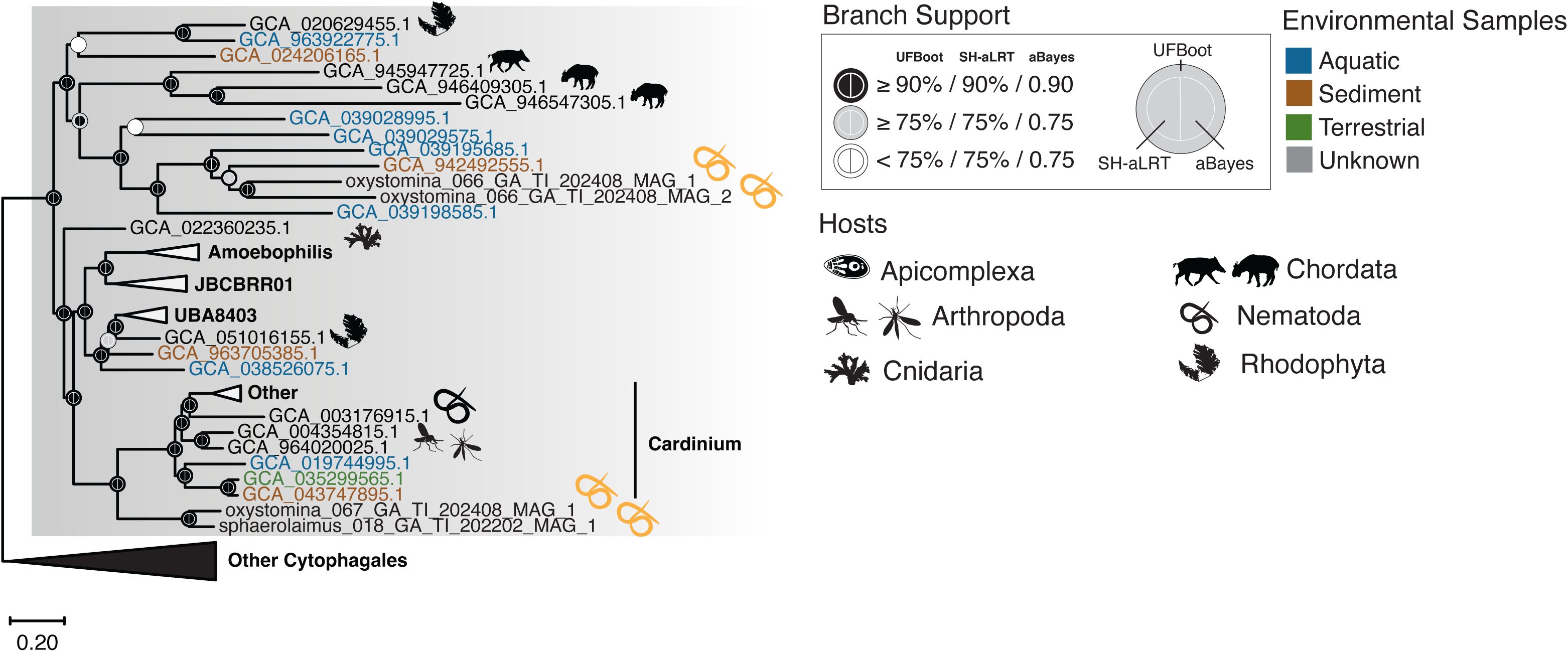
Recovery of nematode-associated *Amoebophilaceae* expands host range of the reproductive manipulator endosymbiont *Cardinium*. A maximum-likelihood phylogenetic tree of *Amoebophilaceae* was generated with 90 single-copy genes (SCGs) using 45 GTDB species-level representatives, 4 *Amoebophilaceae* genomes recovered in this study, and 10 other non-*Amoebophilaceae Cytophagales* genomes to serve as an outgroup were included in the final tree. Phylopics are used to indicate hosts. Golden nematode phylopics indicate nematode-associate MAGs generated from this study. Two nematode-associated *Cardinium* MAGs form a sister-clade with terrestrial invertebrate-associated *Cardinium*. Two additional nematode-associated *Cardinium* MAGs form a clade with *Cardinium* from marine and terrestrial environmental samples.

Finally, we carried out further investigations of the *Patescibacteriota* MAGs found in association with marine nematodes. *Patescibacteriota* is a diverse phylum containing small bacterial epibionts with reduced genomes and this clade is also sometimes referred to the Candidate Phyla Radiation (Méheust et al., 2019; Srinivas et al., 2024). We recovered a surprising number of *Patescibacteriota* MAGs in our nematode dataset (27 MAGs spanning 14 nematode specimens across five marine orders sampled). There is a general paucity of knowledge regarding the natural hosts of most *Patescibacteriota* epibionts, and our study design did not allow us to assess whether *Patescibacteriota* MAGs were nematode symbionts or epibionts living on other bacteria within the nematode holobiont. Nevertheless, to assess the phylogenetic diversity of *Patescibacteriota*, we constructed a phylogeny containing GTDB species-level representatives and 23 *Patescibacteriota* genomes with sufficient SCGs recovered in this study (**Figure S5**). *Patescibacteriota* recovered from marine nematode hosts were not constrained to a single *Patescibacteriota* clade, but rather were found to span the entire phylum. The most common *Patescibacteriota* clades recovered in the metagenomics dataset were *Saccharimonadia* (n = 7), *Dojkabacteria* (n = 4), and JAEDAM01 (n = 4). Notably, three MAGs appeared to represent novel genera with FastAAI values less than 62.30%, while 20 other MAGs represented novel orders and families with FastAAI values 40.42%-52.98%, indicating that nematode holobionts likely harbor deeply divergent phylogenetic lineages within the *Patescibacteriota*.

### Novel chemoautotrophic bacteria (Ca. Thionematobacteraceae) associated with shallow-water and deep-sea nematodes

While exploring our dataset, we observed a total of 11 MAGs classified by the GTDB-Tk as belonging to an unnamed GCF-002020875 bacterial clade. These MAGs were consistently recovered from multiple specimens in two marine nematode families, chemosynthetic Stilbonematidae nematodes (*Eubostrichus sp.*) sampled from tropical coral sands in the Florida Keys, and predatory deep-sea Selachinematidae nematodes collected from sediments in the anoxic Santa Monica Basin off the coast of California. One additional GCF-002020875 bacterial MAG was recovered from another shallow-water nematode specimen from the Florida Keys (Chromadoridae). Phylogenetic analysis using 74 SCGs placed all GCF-002020875 nematode-associated MAGs into a monophyletic clade closely related to the bacterial genus *Thiopontia* (**Figure 5A**). *Thiopontia* are a genera of sulfur-oxidizing bacteria that have been described from anoxic environments in the Black Sea and Cariaco Basin and anchialine ecosystems in Australia (Sun et al., 2023; van Vliet et al., 2021).

**Figure 5.**
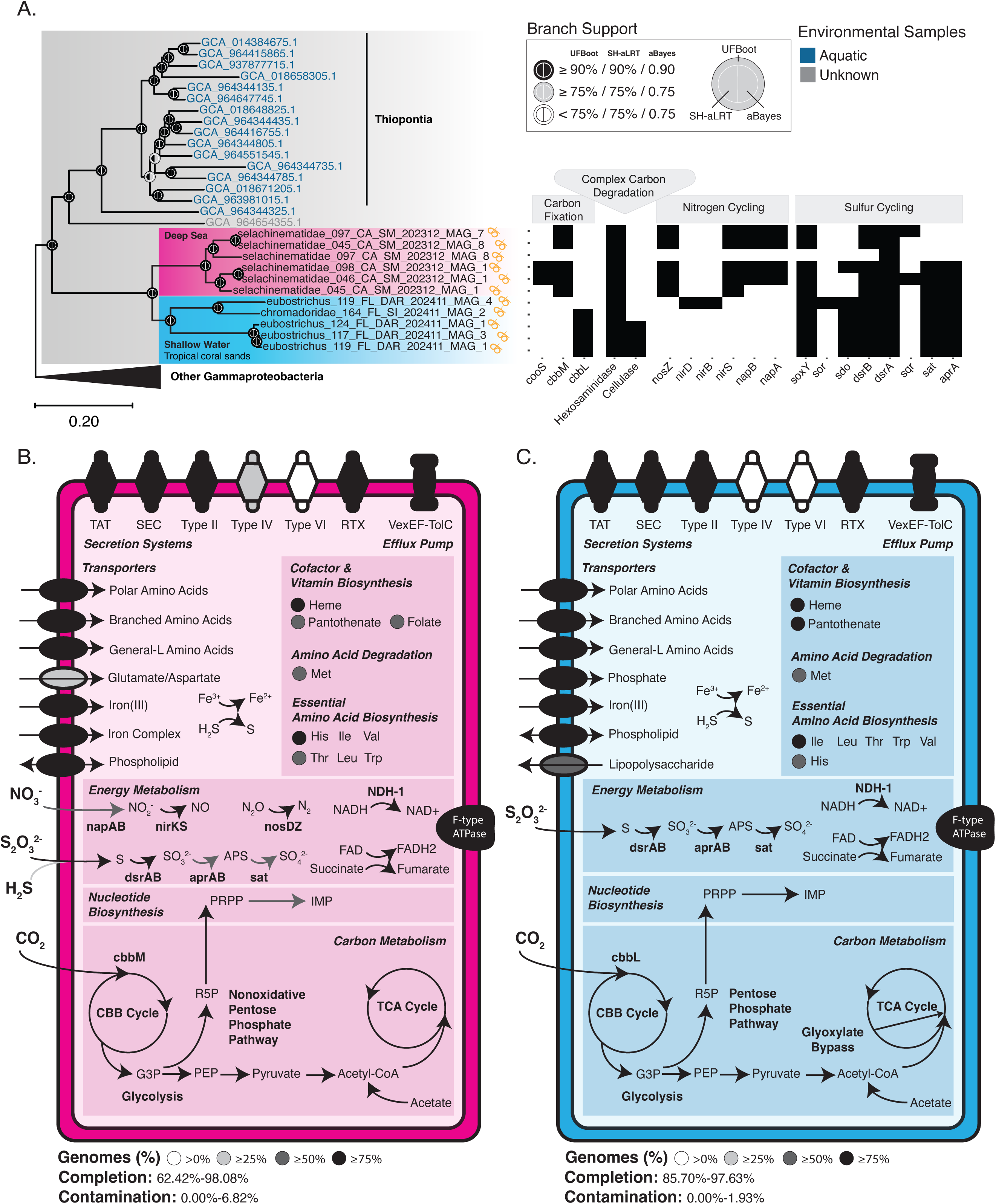
Novel chemoautotrophic bacteria associated with both deep-sea and shallow-water lineages of marine nematodes. A maximum-likelihood phylogenetic tree of *Gammaproteobacteria* was constructed with 74 single-copy genes (SCGs) using 368 order-level representatives, 17 species representatives for the family GCF_002020875, 11 nematode-associated genomes, and 22 *Alphaproteobacteria* genomes to serve as an outgroup. A presence/absence of select genes involved in sulfur and nitrogen cycling, carbon fixation, and carbon degradation were included. The nematode-associated genomes form a sister clade to *Thiopontia* that were sequenced from aquatic environmental samples. Eight of the nematode-associated MAGs contained the *aprA* genes in addition to one of the two forms of RubisCO. Nematode-associated bacteria from Santa Monica Basin (red clade), an historically anoxic environment, contained the cbbM (RubisCO II). Bacteria from Florida Keys (blue clade) contained cbbL (RubisCO I). Additionally, nematodes from shallow-water Florida Keys have key genes involved in nitrite and nitrate reduction. All of the genomes contained hexosaminidases and *dsrA* and *dsrB* (dissimilatory sulfate reductase). Golden nematodes indicate nematode-associated MAGs generated from this study.

Nematode-associated GCF-002020875 MAGs were divided into two distinct phylogenetic subclades according to habitat: one representing anoxic deep-sea sediment habitats (associated with Selachinematidae nematode hosts), and another clade comprising shallow-water tropical coral sands habitats (Stilbonematinae and Chromadoridae nematode hosts). The deep-sea and shallow-water GCF-002020875 lineages represent previously uncharacterized bacterial diversity and are here referred to as *Candidatus* Thionematobacter and *Candidatus* Thiotropica, respectively. Together, these proposed bacterial genera constitute the proposed family *Candidatus* Thionematobacteraceae. Delineation of this family is further supported by FastAAI of 65.82%-90.00% and 55.94%-90.00% within members of the genus *Ca.* Thionematobacter and *Ca.* Thiotropica, respectively. FastAAI between members of different genera are 54.49%-55.48%. FastAAI between members of *Ca.* Thionematobacteraceae and other non-Thionematobacteraceae members of the order GCF-002020875 is <50%. Following SeqCode recommendations (Hedlund et al., 2022), three species names were proposed within this novel chemoautotrophic family (**Table S3**).

Metabolic reconstruction of *Ca.* Thionematobacter genomes revealed that these genomes encode for the core sulfur oxidation pathways, including *dsrAB*, *aprAB*, *sat*, and components of the *Sox* system (**Figure 5BC**), which enable the oxidation of reduced sulfur compounds. In addition, these genomes contain RubisCO, enabling carbon fixation via the Calvin-Benson-Bassham (CBB) cycle. Taken all together, the presence of these metabolic pathways indicated that members of *Ca.* Thionematobacter metabolic capability for sulfur-driven chemoautotrophy. Although all *Ca.* Thionematobacter appear to be novel chemoautotrophs, the two genera exhibit distinct carbon fixation strategies. The deep-sea genus *Ca.* Thionematobacter contains *cbbM* (RubisCO Form II) instead of the *cbbL* gene (RubisCO Form I) found in the shallow-water genus *Ca.* Thiotropica. This divergence in carbon fixation strategies may reflect bacterial evolutionary adaptations to their environmental conditions. RubisCO Form II discriminates less efficiently between oxygen and carbon dioxide compared to RubisCO Form I and may be more advantageous in anoxic environments (Robinson et al., 2003), such as the deep-sea waters of the Santa Monica Basin (**Figure 5**). Furthermore, C*a.* Thionematobacter contain keys genes involved in the denitrification of nitrate (such as *napAB*, *nirKS*, and *nosDZ*), but lacks the gene to reduce nitric oxide to nitrous oxide (*norB*), suggesting an incomplete denitrification pathway (**Figure 5AB**). Phylogenetic analysis of the *aprA* gene suggests that sulfur-oxidation genes within *Ca.* Thionematobacteraceae symbionts have diverse evolutionary origins (**Figure S6**). The bacterial MAG associated with the Chromadoridae nematode contains the *aprA* sulfur-oxidizing gene from *aprA* lineage I, which is closely related to gill sulfur-oxidizing endosymbionts of the coastal bivalve *Solemya velum* (**Figure S6;** (Dmytrenko et al., 2014)). In contrast, *aprA* gene sequences from the remaining *Ca.* Thionematobacteraceae genomes belong to Lineage II and form a monophyletic clade closely related to sulfur-oxiding symbionts of diverse hosts from hydrothermal vents, such as *Tevnia sp.* tubeworms and scaly foot gastropods (Gardebrecht et al., 2012; Nakagawa et al., 2014), and *Ca.* Thiosymbion symbionts from oligochaetes and stilbonematids (Meyer & Kuever, 2007a). Our phylogenetic analysis confirms that the *aprA* genes derive from *Ca*. Thionematobacter MAGs and are unlikely to be the result of contamination from other sulfur-oxidizing chemoautotrophic symbionts, such as *Ca.* Thiosymbion MAGs ectosymbionts attached to *Eubostrichus sp.* nematode hosts. Furthermore, phylogenetic analysis indicates that *aprA* gene of *Ca*. Thionematobacter clusters with *aprA* sequences from other known sulfur-oxidizing bacteria endosymbionts from diverse invertebrate hosts (**Figure S6**; (Watanabe et al., 2016; Zimmermann et al., 2014)).

### Nematode holobionts drive carbon cycling in marine sediments

To assess the metabolic functions of the nematode microbiome and assess their role in nutrient cycling, we used the METABOLIC software package (Zhou et al., 2022) to predict metabolic and biogeochemical traits. Genes involved in complex carbohydrate degradation were specifically subset (**Figure 6B**). The most common carbon degrading pathways recovered from nematode-associated MAGs were hexosaminidase (an enzyme that catalyzes the degradation of chitin; present in ∼65% of MAGs), beta-glucosidase (an enzyme that catalyzes hydrolysis of cellobiose; present in ∼25% of MAGs), and cellulase (enzymes that catalyzes the breakdown of cellulose; present in ∼23% of MAGs). These results indicate that carbon-degrading pathways are functionally conserved in marine nematode holobionts, with phylogenetically diverse nematode hosts maintaining carbon degrading capabilities in their bacterial associates regardless of taxonomy. In terms of sulfur cycling potential in marine nematode holobionts, genes involved in sulfide (*sat*) and sulfate (*sdo*) reduction were prevalent across some bacterial phyla (>20%), including *Acidobacteriota*, *Actinomycetota*, *Chloroflexota*, *Desulfobacterota*, *Pseudomonadota*, and *Thermoproteota* (**Figure 6A**). Key genes that encode the sox complex (*soxBCY*), which mediates the oxidation of thiosulfate to sulfate, were found in low abundances in the *Pseudomonadota* MAGs, while sulfate reduction (*dsrAB)* was found in high abundances in the *Desulfobacterota* MAGs. Furthermore, key genes involved in nitrate reduction (*napAB* and *narHG*) were found in ∼25% and ∼15% in the *Plancomycetota* and *Pseudomonadota* MAGS, respectively. In contrast to the broad conservation of carbon-degrading pathways, genes for sulfur and nitrogen cycling had patchier distributions across nematode-associated MAGs and appeared to be specifically concentrated within certain taxonomic groups.

**Figure 6.**
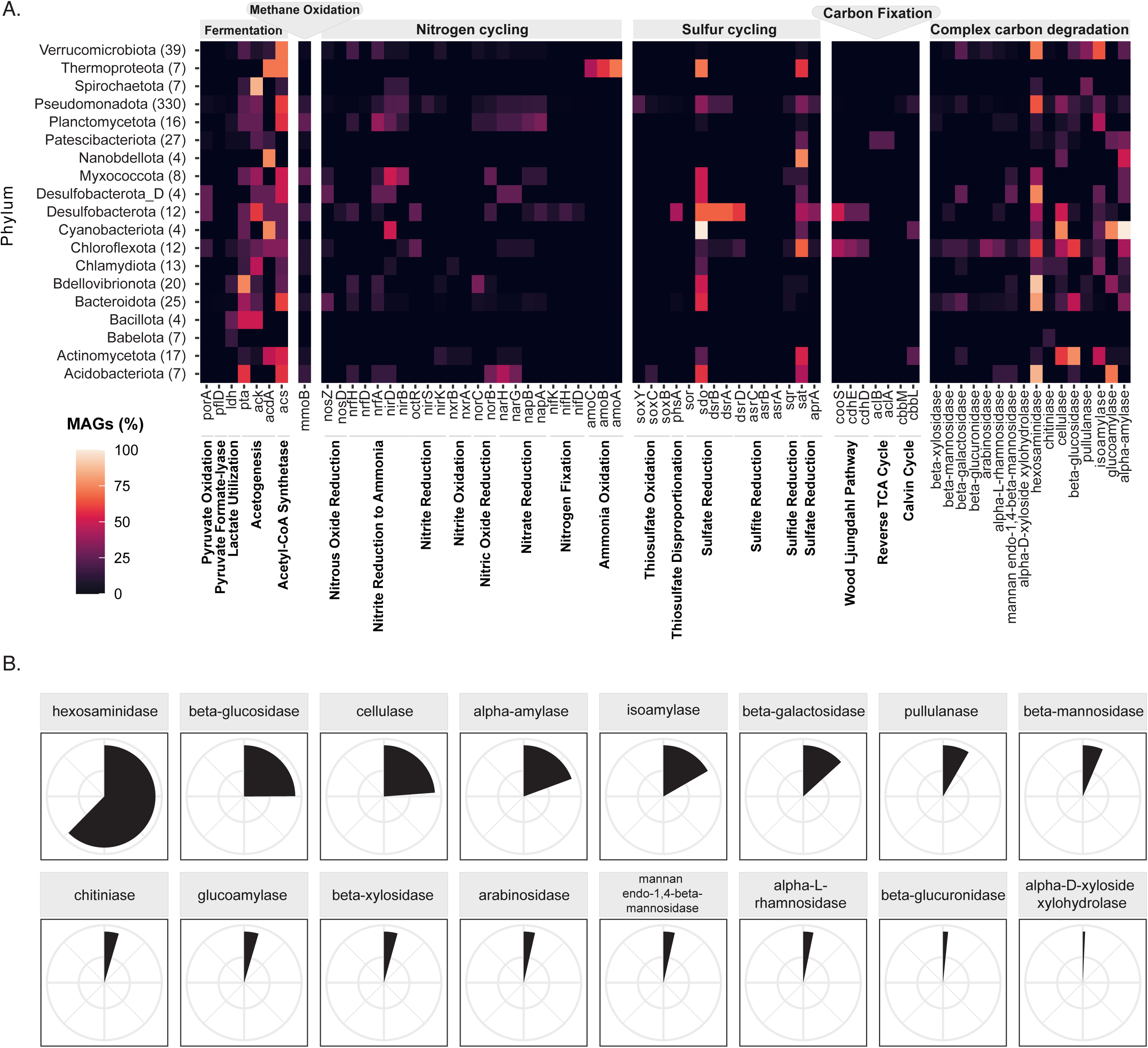
The nematode microbiome underpin key roles in nutrient cycling and degradation of complex carbohydrates. A) Heatmap showing the % of MAGs from each phylum that have select genes/functions. Numbers in parenthesis indicate the total number of MAGs in each phylum. Only phyla with >3 MAGs are shown. B) Percent of all MAGs that contain sixteen enzymes that can degrade complex carbohydrates, ordered from most to least common in nematode-associated holobiont. The most common carbon degrading carbohydrates are hexosaminidase (∼65%), beta-glucosidase (∼25%), and cellulase (∼23%).

### Nematode symbionts occupy distinct physical niches and can be co-located

To assess where nematode-associated microbes are localized within their hosts, we used fluorescent *in-situ* hybridization (FISH) approaches to visualize bacteria using *Pseudoalteromonas*-specific and broad Eubacteria probes in Oncholaimidae (Enoplida) and *Tricoma sp.* (Desmoscolecida) nematodes from Tybee Island, GA and tropical coral sands in the Florida Keys, respectively (**Figure 7 and Figure S7**). In a previous study of Oncholaimidae nematodes (De Santiago et al., 2026), *Pseudoalteromonas* was detected in two distinct host tissues, forming biofilms in the esophagus and ovaries of adult worms. Our work in the present study additionally confirmed that multiple symbionts appear to be co-located within the ovaries of adult female Oncholaimid nematodes, *Pseudoalteromonas* (yellow FISH probe; **Figure 7CDFG**) and another symbiont detected with a general Eubacteria probe (red FISH probe; **Figure 7CDFG**). Similarly, in *Tricoma sp.* specimens representing a different taxonomic order of nematode hosts, we detected both *Pseudoalteromonas* and other co-located symbionts visualized with general Eubacteria probe (**Figure 7FG**). Notably, *Pseudoalteromonas* biofilms were detected in the spermatophores of male *Tricoma sp.* nematodes, and symbiont FISH signals were localized both within nematode tissues (i.e., in reproductive organs) and as external biofilms on the exterior of the nematode (i.e., *Pseudoalteromonas* biofilms strongly concentrated within trough-shaped depressions between desmen rings on the outside of the nematode cuticle; **Figure 7FG**). Our FISH approaches confirm that nematode hosts exhibit multiple distinct physical niches within/outside the host body that can be colonized by symbionts, and co-location of multiple bacterial symbionts appears to be common across phylogenetically diverse nematode hosts.

**Figure 7.**
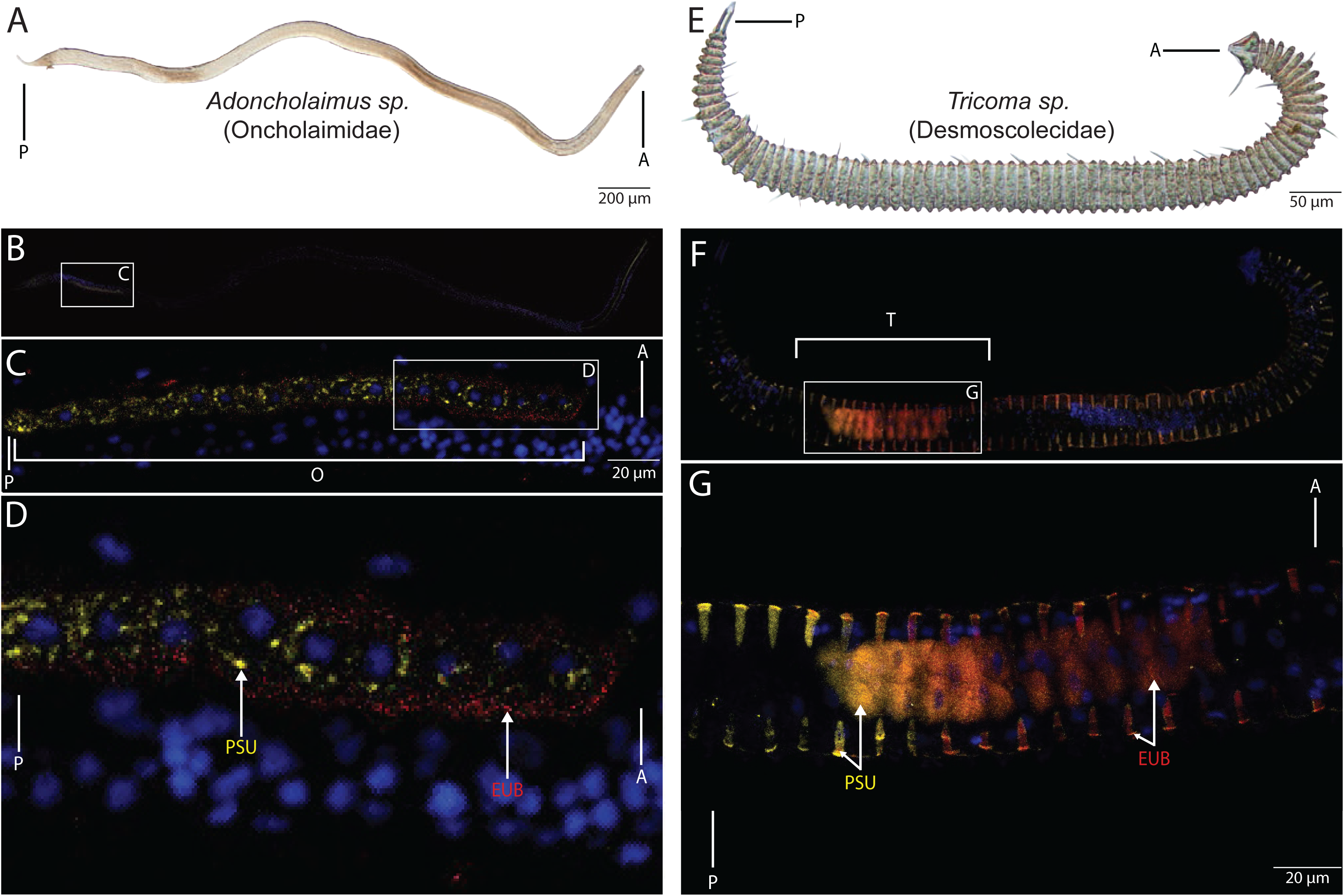
Nematode holobiont taxa occupy distinct physical niches and are often co-localized in reproductive organs. The A) female *Adoncholaimus* (Oncholaimidae) and E) male *Tricoma* (Desmoscolecidae) worms were collected from Tybee Island and Florida Keys, respectively. B-D) *Pseudoalteromonas* (yellow) and other unidentified bacteria (red) in a single underdeveloped ovary of an *Adoncholaimus* (Oncholaimidae) nematode and F-G) the spermatophore of a *Tricoma* (Desmoscolecidae) worm. Micrographs of the Oncholaiomid nematode were adapted from De Santiago et. al. (2026).

## Discussion

Our genome-scale dataset highlights the prevalence of bacterial symbiosis in microscopic marine invertebrates, suggesting broad evolutionary conservation of symbiont lineages within Nematoda and across other animal phyla. Furthermore, the consistent recovery of terrestrial symbiont groups in marine nematode hosts – with marine and terrestrial symbionts often forming sister clades in phylogenetic trees – emphasizes that microscopic marine invertebrates are an underexplored reservoir of host-associated prokaryotic taxa, and suggests that many well-studied terrestrial symbionts may have undiscovered marine relatives. Importantly, our dataset includes the first confirmed report of *Cardinium* endosymbionts from marine invertebrates, a vertically transmitted reproductive manipulator known to infect plant-parasitic nematodes and >800 terrestrial arthropod species (Guo et al., 2026; Mathieson et al., 2025; I. Schön et al., 2025; Tarlachkov et al., 2026). The majority of our marine nematode-associated MAGs were concentrated within just three phyla that are well known for containing other host-associated lineages: *Pseudomonadota*, *Bacteroidota*, and *Verrucomicrobiota*. The *Pseudomonadota* are a diverse bacterial group containing heterotrophic *Gammaproteobacteria* as well as intracellular obligate endosymbionts such as *Rickettsiales* and *Wolbachia (M. E. Schön et al., 2022)*, and bacterial taxa within this phylum are well-studied symbionts of sap-feeding insects, blood-feeding leeches, hydrothermal vent invertebrates, and coastal fish (Brentassi & de la Fuente, 2024; Breusing et al., 2022; Urbanczyk et al., 2012; Zimmermann et al., 2014). Furthermore, recent studies have suggested that *Pseudomonadota* comprise >90% of facultative symbionts in certain insect groups (Palanichamy et al., 2025), and species within this phylum appear to mediate cross-trophic ecological interactions between plants and bactivorous nematodes that serve to suppress pathogens (Chuai et al., 2026). Similarly, the *Bacteroidota* contain keystone taxa associated with the human gut and other animal microbiomes (Comstock & Coyne, 2003; Y. Jiang et al., 2025; Pisaniello et al., 2023; Wexler, 2007; G. Zhu et al., 2022), and many species within this phylum maintain long-term evolutionary relationships with diverse insect hosts (Choi et al., 2025). Likewise, members of the *Verrucomicrobiota* represent key partners ensuring successful initiation of the squid-vibrio symbiosis (McAnulty et al., 2023), appear to produce mandelalide cytotoxins within host tunicate species (Weyna et al., 2015), and species such as *Akkermansia muciniphila* serve as key mucin-degrading species in the human intestinal tract (Chiantera et al., 2023). Notably, we recovered seven Akkermansiaceae MAGs (*Verrucomicrobiota*) from marine nematodes, emphasizing that even some symbiont groups described from humans and vertebrates may be broadly relevant for the ecology and evolution of both marine and terrestrial invertebrates. Taken together, host-associated lineages recovered from marine nematodes are largely consistent with known microbiome associates and obligate symbionts in diverse animal phyla.

At minimum, we estimate that ∼20% of marine nematodes harbor an obligate intracellular symbiont (based on MAG recovery alone), but that estimate may be overly conservative as we recover ∼34% of nematodes hosting obligate symbionts when we extend our analysis to include read-mapping data from single-copy gene windows. This estimate for nematodes is surprisingly consistent with data from tardigrades (reporting endosymbiont prevalence in 9-40% of specimens; (Guidetti et al., 2020)) and arthropods (where 20% of insect species are estimated to host bacterial symbionts; (Cornwallis et al., 2023; Douglas, 2020; Venn et al., 2008). Obligate insect symbioses generally arise in order to solve nutrient deficiencies in host diets (Cornwallis et al., 2023), and endosymbionts such as *Wolbachia* provide B vitamins to their hosts that support insect growth, reproduction, and survival (Douglas, 2017; Nikoh et al., 2014). In nematodes and tardigrades, the reproductive organs appear to be an important physical niche for *Wolbachia*-like endosymbionts (*Rickettsiales*) and other bacterial taxa such as *Pseudoalteromonas* (De Santiago et al., 2026; Guidetti et al., 2020; Hagenbeek et al., 2026), and the present study confirms that male spermatophores as well as female ovaries are both likely hotspots for invertebrate symbioses. Some endosymbionts persist at low frequencies within a population (Hagenbeek et al., 2026) and are known to have a feminizing or male-killing effect in invertebrates (Cordaux et al., 2011; Mathieson et al., 2025), acting as reproductive manipulators while supplementing host nutrition (i.e., B vitamins) to offset the fitness costs of infection (Douglas, 2017). Furthermore, our data from marine nematodes suggests that the coexistence of multiple symbiont lineages within a single host may be more common than previously acknowledged (with 38.27% of marine nematode hosts exhibiting MAGs from multiple obligate intracellular symbiont clades). These data add to the growing recognition that both strain diversity in endosymbionts and multi-species symbioses can provide key benefits to both marine and terrestrial invertebrate hosts (Ansorge et al., 2019; Buysse et al., 2021; Hubert et al., 2025), and the complexity of invertebrate symbioses needs to be considered in light of host/symbiont fitness tradeoffs and population-level evolutionary ecology.

The present study also provides a framework for unifying our knowledge of symbiosis within Phylum Nematoda, one of the most species-rich invertebrate groups on earth (Ingels et al., 2014; Vanreusel et al., 2023). Our holobiont data from marine nematodes – spanning the majority of early-branching clades in Phylum Nematoda (Ahmed & Holovachov, 2021) – establishes important links to model organism research in genetically tractable nematode species such as *Caenorhabditis elegans* and *Pristionchus pacificus* (Nigon & Félix, 2017; Sommer & McGaughran, 2013). Marine nematode holobionts are surprisingly consistent with core microbiome taxa previously reported in a diversity of plant-parasitic and terrestrial nematodes. For example, *Flavobacteriales* appear to be key holobiont associates in nearly all nematode species studied, representing a keystone taxa in the *C.elegans* gut microbiome (Zimmermann et al., 2020), a dominant microbiome taxa that is maintained even under nutrient-starved conditions in *P. pacificus* (Lo et al., 2024), and a bacterial lineage where MAGs were consistently recovered from the holobionts of 25 other terrestrial and plant-parasitic nematode species (Guo et al., 2026). Given that *Flavobacteriales* MAGs were abundantly recovered in six out of eight marine nematode orders in the present study, these patterns suggest strong selection and maintenance of host-associated *Flavobacteriales* over evolutionary timescales in the Nematoda. Similarly, *Verrucomicrobiota* and *Chitinophagales* bacterial MAGs were also consistently recovered in our marine dataset and another large study of terrestrial nematode holobionts (Guo et al., 2026). The evolutionary conservation of these bacterial holobiont taxa suggests that fitness effects are also likely to be conserved across marine and terrestrial nematode hosts, as has been demonstrated recently with experimental work using nematode bacterial isolates (Xue et al., 2024).

At the same time, some nematode holobiont taxa do not seem to exhibit evolutionary conservation, and certain MAGs appear to be restricted to specific nematode lineages. Host identity thus appears to determine at least some symbiont partnerships. For example, phylogenetically divergent Rickettsiales MAGs were recovered from different nematode genera, and when multiple Rickettsiales MAGs were recovered from the same nematode genus they formed a monophyletic clade. *Lariskella* (*Rickettsiales*) MAGs were only recovered from Enoplid nematodes, albeit spanning distant sampling locations in Japan and Georgia, USA. Similarly, Bin98 and novel chemoautotroph *Ca*. Thionematobacteraceae MAGs were mostly recovered in association with Selachimenatidae and Stilbonematidae nematodes, despite these two hosts spanning ecologically disparate deep-sea and tropical coral sand habitats. Although we did not exhaustively sample all taxa or all habitats, our dataset suggests that host identity likely governs invertebrate symbiosis patterns in some capacity, although more genomic data is needed.

Finally, our data suggests that invertebrate holobionts may underpin carbon cycling in benthic marine habitats. Marine nematode holobionts exhibit functional conservation for chitin degradation, via hexosaminidase, and this gene pathway is conserved across ∼65% of host-associated bacterial MAGs irrespective of taxonomy. Studies of cultured bacteria have previously estimated that only 0.4-10% of marine bacteria are able to degrade chitin (Cottrell et al., 1999; Okutani, 1975), and these low oceanic percentages were later validated by constructing genomic libraries of uncultivated free-living and particle-associated bacteria (Cottrell et al., 1999). The maintenance of hexosaminidase pathways in nematode holobionts is in stark contrast to the general absence of this pathway in most free-living marine bacteria, suggesting that this carbon-degrading pathway provides key benefits for invertebrate hosts. Annually, 100 Gigatons (Gt) of chitin sinks to the ocean floor as part of ‘marine snow’ particle flux, however, chitin does not accumulate in high concentrations typically owed to the high efficiency of chitin-degrading pelagic bacteria (Alldredge & Gotschalk, 1990; Cottrell et al., 1999; W.-X. Jiang et al., 2022). Several marine invertebrate taxa, such as sponges and octocorals, have previously been identified as a reservoir for chitin-degrading bacteria and may be source of novel chinolytic enzymes (Meunier et al., 2024; Raimundo et al., 2021). Thus, benthic invertebrate microbiomes – including the holobionts of microscopic phyla such as nematodes, which exhibit high abundance and biomass in marine sediments (Bar-On et al., 2018; van den Hoogen et al., 2019) – may play a critical role in chitin breakdown in the oceans. Furthermore, the functional conservation of beta-glucosidase and cellulases in a quarter of nematode-associated MAGs suggests a much broader role of meiofauna in facilitating benthic carbon cycling worldwide.

Our broad characterization of nematode holobionts allowed us to assess evolutionary patterns in invertebrate-associated taxa, identify previously unrecognized symbiont lineages associated with marine invertebrate hosts, and generate a comprehensive genomic resource of diverse host-associated MAGs that lack representation in existing reference databases. Molecular approaches optimized for low-biomass specimens (e.g. multiple displacement amplification (MDA) reactions carried out on live meiofauna, which generated the best single-specimen datasets in the present study), are critical for generating high-quality data that produce robust sets of holobiont MAGs. Expanded taxon sampling of host-associated assemblages – especially across “neglected” meiofaunal phyla – are needed to determine the broader prevalence and phylogenetic distribution of symbiont taxa across diverse animal hosts and habitats. Here, we primarily focused on known or predicated symbionts clades identified in previous genomic studies. However, deep phylogenomic analysis of divergent MAGs also led to the identification of novel chemoautotrophic symbionts, highlighting the potential for identifying new symbiont taxa that exist outside of many well-characterized endosymbiont clades. Future studies integrating comparative genomics, metabolic profiling, and FISH approaches will be necessary to further explore novel symbionts of marine invertebrates and to provide broader insights into animal-microbe ecological interactions.

## Materials and Methods

### Isolation of marine nematodes for single-worm metagenomics

To assess the diversity of bacterial/archaeal taxa in marine nematode holobionts, we isolated phylogenetically diverse specimens from the Phylum Nematoda (representing eight taxonomic orders) from seven distinct geographic regions (**Figure 1 and Table S1**). Single nematode specimens (one worm per tube) were prepared for short-read metagenomic sequencing, in order to recover a host genome skim alongside the entire holobiont community. A total of 220 marine nematodes were isolated as as part of previous biodiversity surveys (i.e., Bodega Bay, California (Serrano et al., 2026) and the Beaufort Sea, Alaska (Mincks et al., 2021)) and ongoing projects spanning subtidal locations in the Florida Keys, deep-sea sites in Santa Monica Basin off the coast of Southern California, intertidal sites around Tybee Island in Georgia, and deep-sea sites on the Antarctic continental shelf (**Table S1**). Nematodes were isolated from sediments using a decantation-floation method (Danovaro, 2009) and washed over a 45 μm sieve using artificial seawater (Instant Ocean® Spectrum Brands, Inc., USA) that was prepared with Milli-Q ultrapure water. Individual nematode specimens were identified to the lowest possible taxonomic level under light microscopy (typically family or genus level) using the appropriate taxonomic keys for marine nematodes (Platt et al., 1983, 1985; Platt & Warwick, 1988; Warwick et al., 1998) and following current taxonomic classifications on World Register of Marine Species (WoRMS) (WoRMS Editorial Board, 2026). Individual nematodes were quickly rinsed in molecular-grade water to remove transient environmental microbes attached to the cuticle and reduce environmental contaminants before being transferred in microcentrifuge tubes containing either 20 μL of water, 10 μL of REPLI-g Single Cell Cryo-protect Reagent (QIAGEN), or 4 μL of REPLI-g Single Cell Cryo-protect Reagent. Most marine nematodes were identified and transferred into storage buffers while alive. However, nematodes from Bodega Bay were first frozen and stored in a −80 °C freezer before being isolated and stored in microcentrifuge tubes. Nematodes collected from an Antarctic cruise in 2020 were first frozen in DESS at −80°C (Yoder et al., 2006) before being transferred to 10 μL REPLI-g Single Cell Cryo-protect Reagent (QIAGEN).

As described in De Santiago et al. (2026), worms stored in 20 μL of molecular-grade water underwent DNA extraction using the E.Z.N.A MicroElute Genomic DNA kit following the manufacturer’s protocol. Some samples with low DNA yield underwent whole-genome amplification using Genomiphi. Samples stored in 4 μL of REPLI-g Single Cell Cryo-protect Reagent underwent DNA lysis and multiple displacement amplification (MDA) using the REPLI-g Advanced DNA Single Cell Kit (QIAGEN) following the manufacturer’s protocol. Samples in 10μL of REPLI-g Single Cell Cryo-protect Reagent underwent DNA lysis and multiple displacement amplification (MDA) using the REPLI-g Advanced DNA Single Cell Kit (QIAGEN) with a scaled protocol to accommodate increased initial storage volume of the worms, as follows: total of 6 μL of Buffer D2, containing 0.5 μL of 1 M DDT and 5.5 μL of reconstituted buffer DLB was added in the initial step a total of 6 μL of Stop Solution was added after the initial reaction, and amplification reactions were 50 μL total reaction, containing 9 μL of water, 29 μL of REPLI-g advanced sc Reaction Buffer, 2μL of REPLI-g DNA polymerase, and 10 μL of denatured DNA.All RepliG reaction mixtures were incubated in a Thermomixer heated shaker block (Eppendorf, Hamburg, Germany) at 65°C for 10 min, followed by incubation in a T1000 Thermocycler (Bio-Rad) at 30°C for 2 hours with a lid temperature of 70°C, followed by 65°C for 2 min to inactivate the amplification reaction. All reactions were subsequently subjected to magnetic bead cleanup using AMPure XP (Beckman Coulter), and all nematode holobiont samples were subsequently quantified on Qubit 4 Fluorometer (Invitrogen). Single-nematode samples that had sufficient quantities of purified DNA were sent to the Department of Energy Joint Genome Institute, Georgia Genomics and Bioinformatics Core at the University of Georgia, or the Roy J. Carver Biotechnology Center at the University of Illinois at Urbana-Champaign for Illumina library preparation and Illumina sequencing using the NextSeq or NovaSeq Platform (**Table S1**).

### Metagenome assembly and extraction of host DNA barcodes

Illumina sequences were processed using MeioBIOME, a snakemake workflow developed for the parallel analysis of host genome skims and any host-associated bacterial/archaeal genomes recovered from single-specimen metagenomic datasets (De Santiago & Bik, 2026). As the first step in the meioBIOME workflow, read quality and adapter content were summarized using FastQC (Andrews & Others, 2010) and MultiQC (Ewels et al., 2016). Reads underwent adapter and quality-trimming using Trimmomatic (Bolger et al., 2014), with a leading and trailing PHRED score of 2 and a SLIDINGWINDOW:4:20. Only reads with a minimum length of 55 bps, the maximum kmer length implemented by metaSPAdes (Nurk et al., 2017) following default parameters, were kept for assembly and downstream analysis. To reduce the computational resources required for metagenome assembly, samples were deduplicated with fastp (Chen et al., 2018) using default parameters (deduplication accuracy of 3, see fastp manual for more details). Quality-controlled samples were then assembled using metaSPAdes (Nurk et al., 2017), and scaffolds shorter than 1,000 bp were filtered from the dataset. To confirm the taxonomic identification of each nematode specimen (and identify any other eukaryotes present in the nematode holobiont), the 18S rRNA gene was extracted using two methods. First, phyloFlash (Gruber-Vodicka et al., 2020) was used on raw data to extract reads that successfully mapped to the SILVA reference databases before assembling them with metaSPAdes. Additionally, 18S rRNA genes were extracted from the metagenome-assembled scaffolds using Barrnap (https://github.com/tseemann/barrnap). The nematode 18S rRNA sequences were queried against the NCBI nucleotide database using BLAST+ (Camacho et al., 2009) to confirm the family or genus-level identification of the host.

### Binning host-associated bacterial and archaeal MAGs

Host-associated bacteria/archaea were analyzed by binning metagenome-assembled genomes (MAGs) using three binning algorithms, and MAGs were further optimized and dereplicated using DASTool (Sieber et al., 2018). First, each nematode holobiont sample was binned using CompleteBin (Zou et al., 2025), COMEBin (Wang et al., 2024), and MetaBat2 (Kang et al., 2019). CompleteBin and COMEBin are two state-of-the-art binning tools that implement a dynamic contrastive learning approach, and both have been shown to improve the number of recovered MAGs in real-world metagenomic datasets and in simulated datasets with high strain-level variability. DASTool (Sieber et al., 2018) was run using a score threshold of 0 to output an optimized set of dereplicated (non-redundant) MAGs for each metagenomic assembly without filtering for quality (as determined by DASTool). Following recommended microbial community standards (Bowers et al., 2017), a custom Python script was used to keep bins with a minimum 50% of completion and less than 10% estimated contamination (according to a set of bacterial and archaeal single-copy genes employed by DASTool). Next, MAGpurify (Nayfach et al., 2019) was implemented on each MAGs using four internal modules to remove contigs that were misbinned. The *known-contam* modules were used to remove human and phi-x contaminants, and the *gc-content* module was used to remove contigs with outlier GC composition. The *phylo-markers* and *clade-markers* modules were used to remove taxonomically discordant contigs within each MAG. Finally, CheckM2 (Chklovski et al., 2023), which uses various genomic features, such as amino acid count and number of coding sequences, in addition to single-copy genes, was run on the taxonomically concordant MAGs to reassess quality. The GTDB-tk workflow (Chaumeil et al., 2022), while skipping the ANI screen, was used to assign taxonomy to each MAG. A total of 815 MAGs were recovered from the 215 single-specimen nematode metagenome samples. Of the 815 MAGs, 270 were designated as high-quality, 381 as medium-quality, and 164 as low-quality (**Table S2**). We used a custom script to estimate genome size for each MAG following the equation:

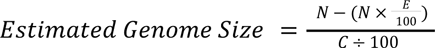

where, ‘N’ is the number of nucleotides in each MAG and where ‘E’ and ‘C’ are the percent contamination (i.e., erroneous nucleotides mistakenly binned) and completion, respectively, as estimated by CheckM2. First, the number of contaminant nucleotides was calculated and removed from the total nucleotides in each MAG. Next the total number of “non-contaminant” nucleotides was divided by the estimated completion to estimate the size of the “complete” genome.

### Phylogenetic analysis of nematode holobionts and endosymbiont lineages

To assess the overall microbial biodiversity of marine nematode holobionts, symbiont clades were placed into a phylogenomic framework using GTDB representative genomes downloaded using GToTree (M. D. Lee, 2019). Clade-specific phylogenies were generated for a subset of the symbiont-exclusive clades (i.e., *Rickettsiales*, *Chlamydiota*, Bin98, and *Patescibacteriota*) using the GToTree workflow (M. D. Lee, 2019). Symbiont-exclusive clades are defined as bacterial lineages where ≥90% of all members were previously predicted to be obligate intracellular symbionts. Symbionts are either known symbionts that are comprehensively described in the literature and newly predicted symbiont lineages based on genome characteristics (Villada et al. 2025). The previously described symbiont-exclusive taxonomic clades were used to assign lifestyle to MAGs recovered from the nematode dataset. GToTree HMMs containing bacterial and phyla-specific SCGs were used to generate symbiont phylogenies. For the *Rickettsiales* phylogeny, 74 bacterial SCGs were extracted from 650 GTDB species representatives, 20 nematode-associated MAGs were included from this study, and 25 *Gammaproteobacteria* were included as an outgroup. For the *Chlamydiota* phylogeny, 74 bacterial SCGs were extracted from 560 GTDB species representatives, 12 nematode-associated MAGs were included from this study, and 20 non-*Chlamydiota* representative genomes from the PVC superphylum (10 *Verrucomicrobiota* and 10 *Planctomycetota*) were included to serve as an outgroup. For the Patescibacteriota, 25 prokaryote SCGs were extracted from 845 GTDB family representatives, 27 nematode-associated MAGs were included from this study, and 25 representative genomes from Chloroflexota were used as an outgroup. Due to paucity of Bin98 genomes available in GTDB, a phylogenetic tree was constructed using 74 SCGs with 429 Alphaproteobacteria family representatives (including two Bin98 genomes), 11 Bin98 nematode-associated MAGs derived from the present study, and 25 Gammaproteobacteria family representatives used as an outgroup. For each group, the SCGs were aligned by synteny and masked using the GToTree workflow following default parameters. Genome with low recovery of SCGs were removed from the dataset. The phylogenomic trees for each clade were then constructed in IQTree using a partition model (Nguyen et al., 2015), which implements ModelFinder (Kalyaanamoorthy et al., 2017) to identify the best protein model for each SCG using Bayesian Information Criteria (BIC). An initial 1,000 parsimony trees were generated and the top 20 initial trees and perturbation strength of 0.20 were used for the initial tree search algorithm. Additionally, the number of unsuccessful iterations to stop was increased from 100 (default) to 500 to allow for better optimization during tree search and prevent premature termination of tree search before converging on an optimal tree topology. Branch support was calculated using three methods: 1) Ultrafast Bootstrap (UFBoot) using 1,000 replicates, 2) SH-like approximation livelihood ratio test (SH-aLRT) with 1,000 replicates, and 3) approximate Bayes test (aBayes). Trees were analyzed using TreeViewer and stylized in Adobe Illustrator. Phylopics were used to clarify host taxonomy: Apicomplexa (Levi Simons), Arthropoda (Carter Johnson, Graham Montgomery, Melissa Broussard, Michael Day, Nathan Jay Baker), Ciliophora (Levi Simons), Cnidaria (Guillaume Dera), Chordata (Mecedes Yrayzoz, Servien, Ferran Sayol, and Kay Caspar), Echinodermatada (Guillaume Dera), Mollusca (Tauana Cunha), Nematoda (Michelle Site), Porifera (Marina Vingiani), Rhodophyta (Arcadia Sciences), Streptophyta (Ricarda Pätsch), Tunicata (Apokryltaros), Xenoceolomorpha (Meyer-Wachsmuth I, Curini Galletti M, Jondelius U (Vectorization by Y. Wong))

To further assess the phylogenetic novelty of the recovered MAGs, we calculated FastAAI between each MAG and its closest GTDB representative genome (Gerhardt et al., 2025). First all GTDB species-representative genomes were downloaded and FastAAI was used to perform pairwise comparison between each recovered MAG and each GTDB genome. Following the recommendations set by the developers, genus- and family- novelty were assessed using the proposed FastAAI thresholds of 62.30% and 53.00%, respectively. Finally, we quantified the abundance and prevalence of symbiont-exclusive clades across all marine nematode holobionts in our dataset. To estimate the abundance of symbiont-exclusive clades, two methods were used. First, SingleM (Woodcroft et al., 2025) was run on raw metagenomic reads, as recommended by the developers, to estimate the taxonomic abundance of bacterial taxa using single-copy genes. The abundance of symbiont-exclusive clades was further visualized by subsetting the SingleM dataset according to the symbiont-exclusive clades described in Villada et al. (2025).

### Assessing the functional potential of the nematode holobiont

To assess the functional genes and metabolic pathways present in marine nematode holobionts, we analyzed the MAGs using METABOLIC-G v4.0 (Zhou et al., 2022). METABOLIC-G was run using prodigal (Hyatt et al., 2010) in metagenomics mode and with a module-cutoff of 0.75. HMMs included in METBOLIC were used to identify genes involved in biogeochemical cycles, fermentation, and complex carbon degradation. Phyla with less than three MAGs were excluded from the analysis. Heatmaps showing the presence/absesnce of select biogeochemical genes were constructed using Ggtree (Xu et al., 2022) in RStudio.

### SeqCode section subheading for Ca. Nematobacteraceae

To formally name the uncultivated chemoautotrophic bacteria GCF-002020875, we followed the SeqCode recommended nomenclature framework for (Hedlund et al., 2022). First, a maximum-likelihood phylogenetic tree of *Gammaproteobacteria* was constructed with 74 single-copy genes (SCGs) using 368 order-level representatives, 17 species representatives for the family GCF_002020875, 11 nematode-associated genomes, and 22 *Alphaproteobacteria* genomes to serve as an outgroup, following the parameters used to generate symbiont phylogenetic trees. The SCGs were aligned by synteny and masked using the GToTree workflow following default parameters. The phylogenomic trees were then constructed in IQTree using a partition model (Nguyen et al., 2015), implementing ModelFinder (Kalyaanamoorthy et al., 2017) to identify the best protein model for each SCG using Bayesian Information Criteria (BIC). An initial 1,000 parsimony trees were generated and the top 20 initial trees and perturbation strength of 0.20 were used for the initial tree search algorithm. Additionally, the number of unsuccessful iterations to stop was increased from 100 (default) to 500. Branch support was calculated using three methods: 1) Ultrafast Bootstrap (UFBoot) using 1,000 replicates, 2) SH-like approximation livelihood ratio test (SH-aLRT) with 1,000 replicates, and 3) approximate Bayes test (aBayes) . Second, we compared FastAAI and FastANI to closely related representative genomes, as determined by their phylogenetic placement (see below). Second, the high and medium-quality MAGs were compared to each other using fastANI via dRep to determine whether recovered nematode-associated MAGs belong to different species clusters. Adhering to SeqCode and recommendations, only species clusters with more than one MAG recovered from different samples were formally described (Hedlund et al., 2022). Following community standards, MAGs with the highest completion (>90%) and lowest contamination (<5%) were selected as the type genome for the proposed taxon (**Table S3**; (Bowers et al., 2017)). FastAAI was calculated to assess whether genome divergence of recovered MAGs supports the proposal of multiple novel genera, using the recommended FastAAI threshold of 62.30% (Gerhardt et al., 2025). MAGs have FastAAI that fall below the family-designation threshold of 53.00% were designated as a novel family.

### Phylogenetic analysis of the aprA gene from the novel chemoautotrophic bacteria family Ca. *Nematobacteraceae*

To assess evolutionary patterns in a novel chemoautotrophic Gammaproteobacteria found in association with both shallow-water and deep-sea marine nematodes, we explored the phylogenetic placement of the adenosine-5′-phosphosulfate reductase alpha subunit gene (*aprA*), which encodes a key enzyme in microbial sulfate reduction and sulfur oxidation (Meyer & Kuever, 2007b). An *aprA* gene tree was constructed using 320 *aprA* sequences that were used in a previous study to construct a high-quality *aprA* gene tree (Watanabe et al., 2016), seven *aprA* genes recovered in this study from nematode-associated MAGs corresponding to *Ca.* Nematobacteraceae chemoautotrophic symbionts, and 27 *aprA* genes from *Ca.* Thiosymbion MAGs that were also recovered from this dataset. The *aprA* genes were aligned using MAFFT with the standard FFT-NS-i algorithm (Katoh & Standley, 2013). The alignment was masked using AliFilter (Bianchini et al., 2026) and the final masked alignment had 644 comparable amino acid positions. A maximum-likelihood gene tree was constructed using IQTree3 (Nguyen et al., 2015), which implements ModelFinder (Kalyaanamoorthy et al., 2017) to identify the best protein model. The best-fit model LG+F+R7 was chosen according to Bayesian Information Criteria (BIC). An initial 1,000 parsimony trees were generated and the top 20 initial trees were used for the initial tree search algorithm. Additionally, the number of unsuccessful iterations to stop was increased from 100 (default) to 200. Branch support was calculated using three methods Ultrafast Bootstrap (UFBoot) using 10,000 replicates. Trees were analyzed using TreeViewer and stylized in Adobe Illustrator.

### Visual confirmation of marine nematode symbionts using FISH

Live marine nematodes from the Florida Keys and Tybee Island, GA, were subjected to fluorescence *in situ* hybridization (FISH) using a modified protocol previously used for visualization of nematode-associated bacteria (Bellec et al., 2019). Live nematodes were fixed for 2 h in 3% formaldehyde with sterile artificial seawater (Instant Ocean Spectrum Brands, Inc., USA), followed by three rinses in a 1:1 solution of Phosphate-Buffered Saline (PBS):sterile seawater. Fixed samples were subsequently transferred to a 1:1 storage buffer (100% ethanol:2x PBS) and stored at −20°C. For FISH, marine nematodes were rinsed in 30% formamide buffer and hybridized in 150 μl hybridization buffer [0.9 M NaCl, 0.02 M Tris-HCl (pH 8.0), 0.01% SDS, 2 ml blocking reagent [5% bovine serum albumin (BSA in 0.5% PBS-Tween 0.5%) and 95% 7.4 pH PBS], and 30% formamide] at 46°C for 3 h with 1 μM concentration of each probe. We used a combination of universal bacterial probes that have been shown to increase coverage of bacterial phyla (EUB338I: 5′-GCTGCCTCCCGTAGGAGT-3′; CY-3; (Amann et al., 1990); EUB338II: 5′-GCAGCCACCCGTAGGTGT-3′; CY-3; EUB338III: 5′-ACACCUACGGGUGGCAGC-3′; CY-3; (Amann et al., 1990; Daims et al., 1999)), alongside a *Pseudoalteromonas*-specific probe (5′-TTGACCCAGGTGGCTGCC-3′; CY-5; (De Santiago et al., 2026; Greuter et al., 2016)), and nonsense probe (5′-ACTCCTACGGGAGGCAGC-3′; FAM; (Wallner et al., 1993)). After probe hybridization, nematodes were rinsed three times in washing buffer [0.102 M NaCl, 0.02 M Tris-HCl (pH 80), 0.005 M Ethylenediaminetetraacetic acid (EDTA), 0.01% SDS] for 30 min at 46°C for a total of 1.5 h. The hybridized specimens were mounted on a slide in ProLong™ Diamond Antifade Mountant with DAPI containing 4′,6-diamidino-2-phenylindole (DAPI) and observed using the Zeiss LSM 880 Confocal Microscope system. The Zen 2.3 imaging software was used for image acquisition and processing. The images of the Oncholaimid nematode were acquired as single-plane confocal images using a 40× oil-immersion objective, whereas the images of the *Tricoma sp.* nematode were acquired as a maximum-intensity projection of a z-stack of single-plane confocal images using a 40X oil-immersion objective. The whole nematode image was generated as tiled acquisitions using identical imaging parameters. All images were subsequently processed using FIJI (ImageJ).

## Acknowledgements

Funding for this study was provided by the Gordon and Betty Moore Foundation (Symbiosis in Aquatic Systems Initiative, grant #9326), and a National Science Foundation CAREER award (DEB-2144304) and Antarctic Program award (OPP-2132641) to HMB at UGA. Research support for ADS was provided by the University of Georgia Research Foundation and the National Institute of General Medical Sciences of the National Institute of Health under award number 1T32GM142623. We acknowledge the Joint Genome Institute (JGI, Proposal ID: 505025), the Georgia Genomics and Bioinformatics Core (GGBC, UGA, RRID:SCR_010994), and the Roy J. Carver Biotechnology Center at the University of Illinois at Urbana-Champaign for metagenomic sequencing of single-worm isolates and the Georgia Advanced Computing Resource Center (GACRC) at UGA (https://gacrc.uga.edu/) for computational resources that have contributed to the results in this publication. The work (proposal: 10.46936/10.25585/60001240) conducted by the U.S. Department of Energy Joint Genome Institute (https://ror.org/04xm1d337), a DOE Office of Science User Facility, is supported by the Office of Science of the U.S. Department of Energy operated under Contract No. DE-AC02-05CH11231. We also thank Kevin Kocot for providing nematode samples from the Antarctic cruise NBP20-10.

## Data Availability

All the genomes used in this study are publicly available in the NCBI RefSeq (see **Supplementary Table S1** for all associated accession codes and references). Raw metagenomic sequences generated by the Georgia Genomics and Bioinformatics Core (GGBC) and the Joint Genomic Institute (JGI, Proposal ID: 505025) were deposited in the NCBI Sequence Read Archive (BioProject: XXXXXX and XXXXXX). We follow the STREAMS guidelines for reporting metadata from environmental and host-associated microbiome studies (Kelliher et al., 2025).

## Code Availability

All scripts and files required for the reproducibility of all analyses conducted in this study are available via GitHub (https://github.com/BikLab/nematode-MAGs).

## Supplementary Figure Legends

**Figure S1. Symbiont-exclusive clades were captured across all well-sampled nematode lineages.** A) The number of specimens where we recovered (Yes; n = 40) and did not recovered non-host 18S rRNA (No; 160) were plotted. N/A indicates samples with failed assemblies (n = 20). 18S rRNA sequences on the right organized by taxonomy. The sum of 18S sequences are >40 since some samples had more than 1 non-host 18S rRNA sequence. B) Cladogram of major nematode lineages, numbers within parenthesis indicate total specimens in each order. Colors of micrographs indicate nematode lineages of select representative specimens. Heatmap indicates the prevalence (%) of symbionts-exclusive clades found across major nematode lineages.

**Figure S2. Symbiont-exclusive taxa are highly prevalent across marine nematodes and coexist in single individuals.** SingleM (Woodcroft et al., 2025) was used to assign taxonomy to the microbiome of each nematode samples and calculate the relative abundance of A) each phyla and B) symbiont-exclusive clades (Villada et al., 2025). Of the 220 nematode holobionts sequenced, 81 had a symbiont (36.82%). Of the samples with symbionts, 31 (38.27%) nematodes had more than one symbiont, wheres 14 (17.28%) worms had more than two symbionts. The five common symbionts were *Patescibacteriota* (44; 20%), *Rickettsiales* (31; 14.09%), *Chlamydiota* (25; 11.36%), *Legionellales* (7; 3.18%), and *Coxiellales* (6; 2.73%). When excluding Patescibacteriota (which are primarily epibionts of bacteria), 56 individuals (25.45%) had a symbiont.

**Figure S3. Novel microbial lineages are recovered across diverse nematode hosts.** FastAAI (Gerhardt et al., 2025) was compared between bacterial MAGs generated in this study and the closest GTDB reference genome. MAGs from four of the most well-sampled orders were compared against GTDB reference genomes: A) Enoplida, B) Desmoscolecida, C) Desmodorida, D) Chromadorida and families. MAGs from four of the most well-sampled families were compared against GTDB reference genomes: E) Oncholaimidae, F) Desmoscolecidae, G) Stilbonematinae (Desmodoridae), H) Selachinematidae.

**Figure S4. Recovered nematode-associated *Chlamydiales* genomes are closely-related to diverse aquatic and sediment samples.** A maximum-likelihood phylogenetic tree of *Chlamydiales* was generated with 74 single-copy genes (SCGs) using 533 GTDB species-level representatives, 7 *Chlamydiales* genomes recovered in this study, and 19 non-*Chlamydiota* representative genomes from the PVC superphylum (10 *Planctomycetota* and 9 *Verrucomicrobiota*) to serve as an outgroup were included in the final tree. Focal subtrees of A) JAAKR01, B) JAJFMA01, C) SM23-29, D) and *Simkaniaceae* were generated by pruning each family from the full phylogenetic tree, with the remaining order shown as a sister reference clade. The non-*Chlamydiota PVC* outgroup is not shown. Phylopics are used to indicate hosts. Golden nematode phylopics indicate nematode-associate MAGs generated from this study.

**Figure S5. Nematode-associated *Patescibacteriota* encompass diverse lineages across the phylum.** A maximum-likelihood phylogenetic tree of *Patescibacteriota* was generated with 25 single-copy genes (SCGs) using 826 GTDB family-level representatives, 22 *Patescibacteriota* genomes recovered in this study, and 25 other *Chloroflexota* genomes to serve as an outgroup were included in the final tree. Golden nematode phylopics and branches indicate nematode-associate bacterial MAGs generated from this study.

Figure S6. *AprA* genes from novel chemoautotrophic symbionts cluster with *aprA* genes from hydrothermal vent sulfur-oxidizing symbiont taxa. An *aprA* gene tree was constructed using the 320 *aprA* sequences from a previous study (Watanabe et al., 2016), 7 *aprA* genes recovered from the chemoautotrophic *Ca.* Nematobacteraceae MAGs, and *aprA* genes from *Ca.* Thiosymbion MAGs that were also recovered from this dataset. A) The *aprA* gene from the Chromadoridae nematode clustered with aprA Lineage I, and B) the rest fo the *Ca.* Nematobacteraceae *aprA* genes were identified as aprA lineage II and *f*ollows similar evolutionary history to *aprA* gene from sulfur-oxidizing chemoautotrophic symbionts of hydrothermal vent taxa, such as tubeworms and vent snails. The *aprA* I and *aprA* II genes from stilbonematid-associated *Ca.* Thiosymbion genomes formed their own phylogenetic clades.

**Figure S7. Fluorescent in situ hybridization of the holobiont of Oncholaimidae and Desmoscolecidae nematodes.** The A) female *Adoncholaimus* (Oncholaimidae) and E) male *Tricoma* (Desmoscolecidae) worms were collected from Tybee Island and Florida Keys, respectively. *Pseudoalteromonas* (yellow) and other unidentified bacteria (red) were found in the reproductive organs of B) a female *Adoncholaimus* (Oncholaimidae) nematode and E) a male *Tricoma* (Desmoscolecidae) worm. Eubacterial probes are stained red (B,F,J,N), whereas *Pseudoalteromonas*-specific probes are yellow (A,E,I,M). Green indicates nonsense probes (C,G,K,O). The nonsense probes were staining the Oncholaimidae host cells, but none of the bacterial cells (C,G).

## Supplemental Table Legends

**Table S1. Summary of the nematodes that were included in this study.** A total of 220 specimens spanning 55 unique genera within 30 families and 8 orders were included in this study. Metadata includes the year the specimen was collected, location, latitude and longitude. Each worm was identified to the family or genus level using nematode taxonomic keys. When possible, taxonomic identification was confirmed using the 18S rRNA gene (see methods).

**Table S2. Summary of the metagenome-assembled genomes (MAGs) that were recovered from single worm holobionts.** total of 220 specimens spanning 55 unique genera within 30 families and 8 orders were included in this study. Metadata includes the year the specimen was collected, location, latitude and longitude. Each worm was identified to the family or genus level using nematode taxonomic keys. When possible, taxonomic identification was confirmed using the 18S rRNA gene (see methods).

Supplementary Table CC. Proposed taxon names of new nematode-associated chemoautotrophic bacteria family Ca. *Thionematobacteraceae*. The proposed taxon ranks following SeqCode recommendations.

