## Supplementary figures and images for "Marine nematodes exhibit widespread symbiosis, novel chemoautotrophy, and evolutionary conservation of holobiont taxa"

### Figure S1

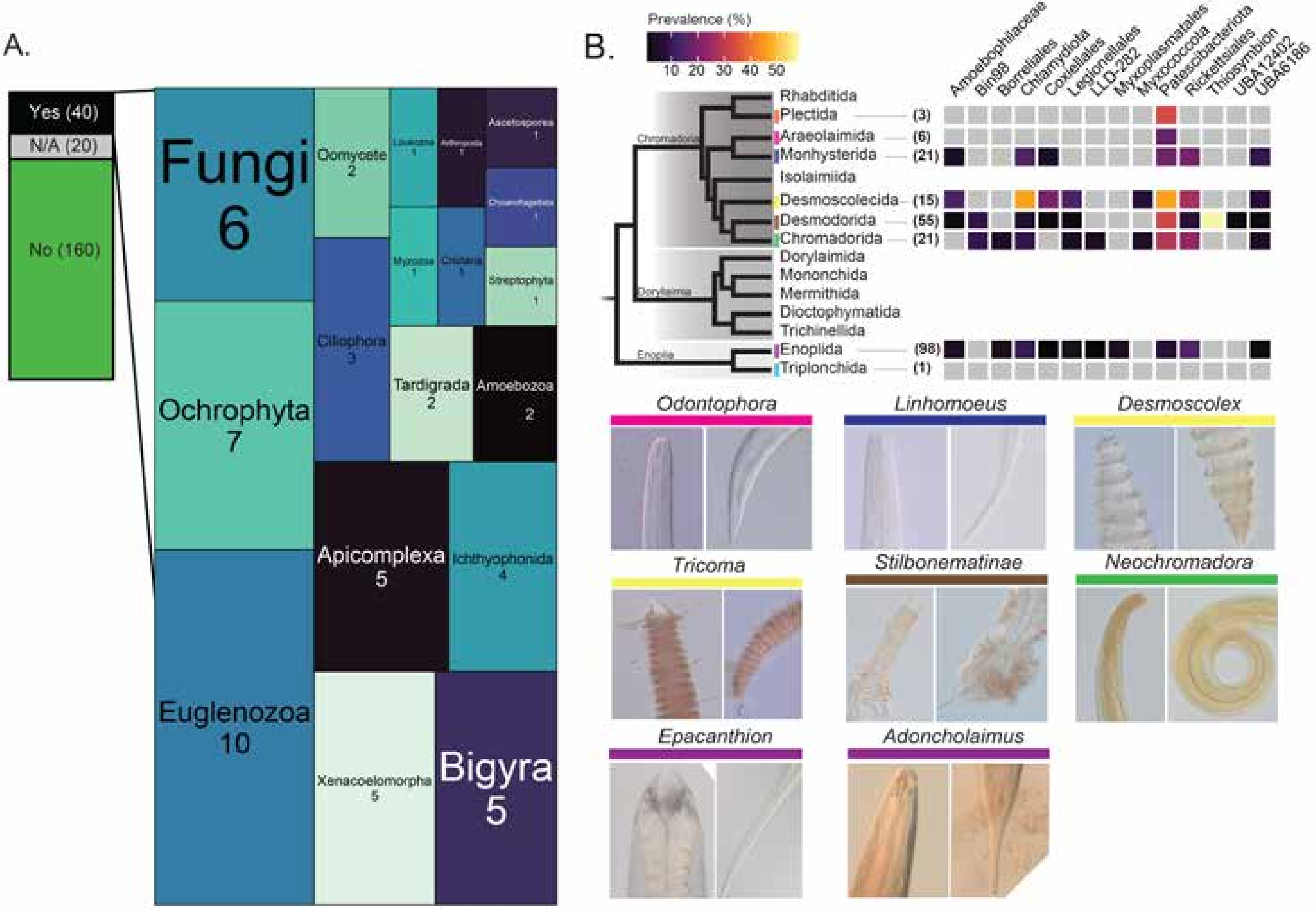

### Figure S2

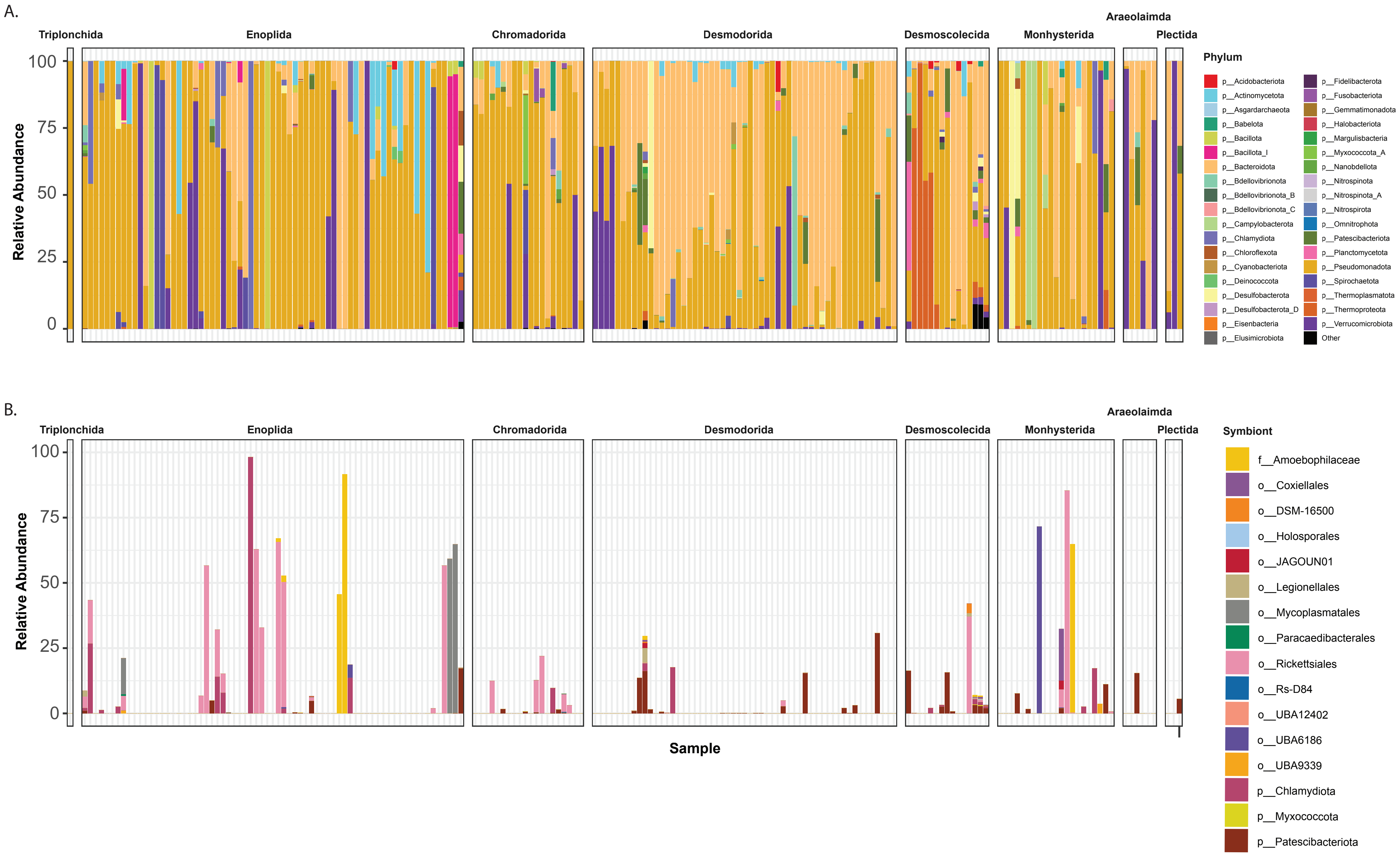

### Figure S3

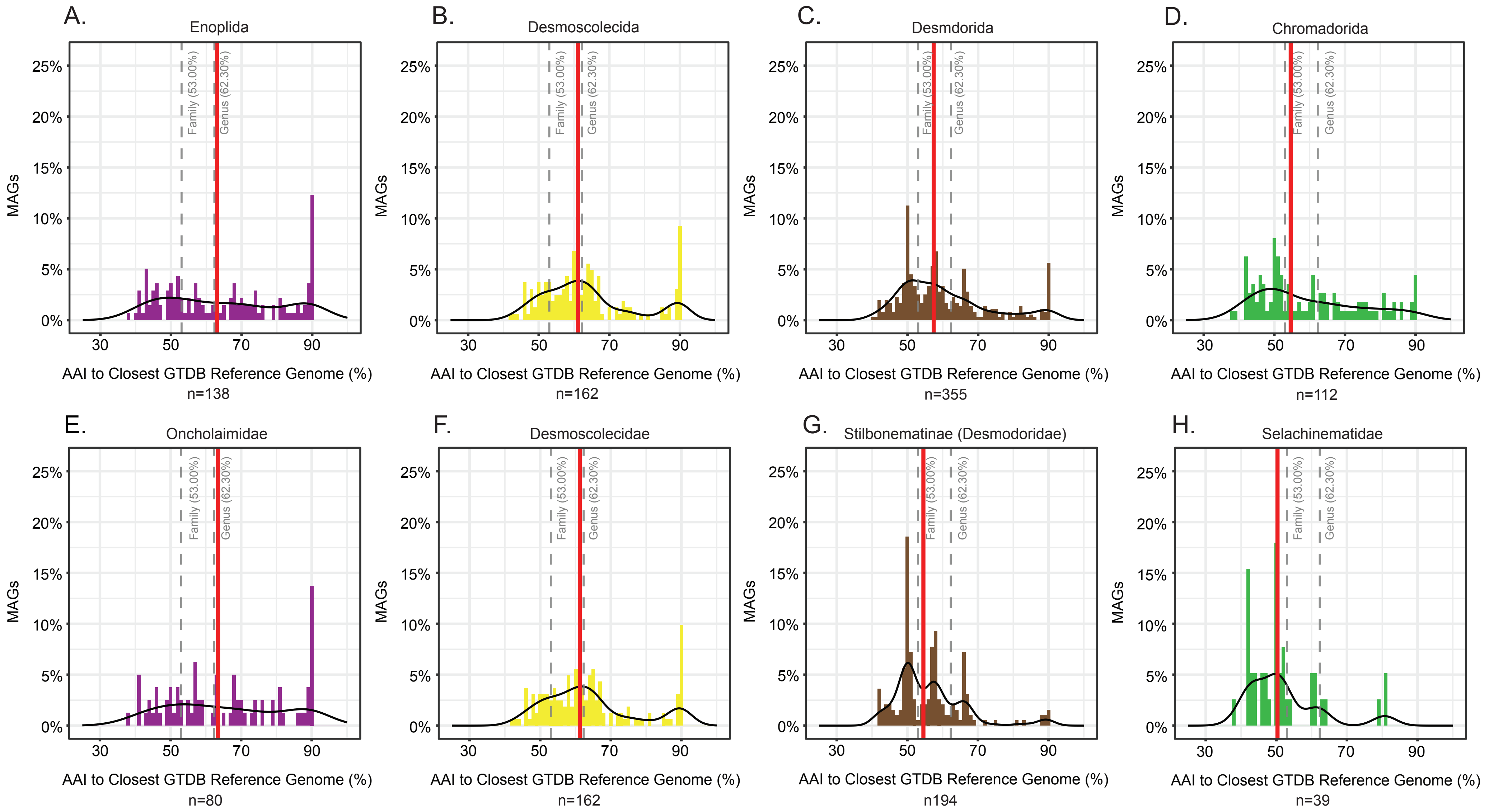

### Figure S4

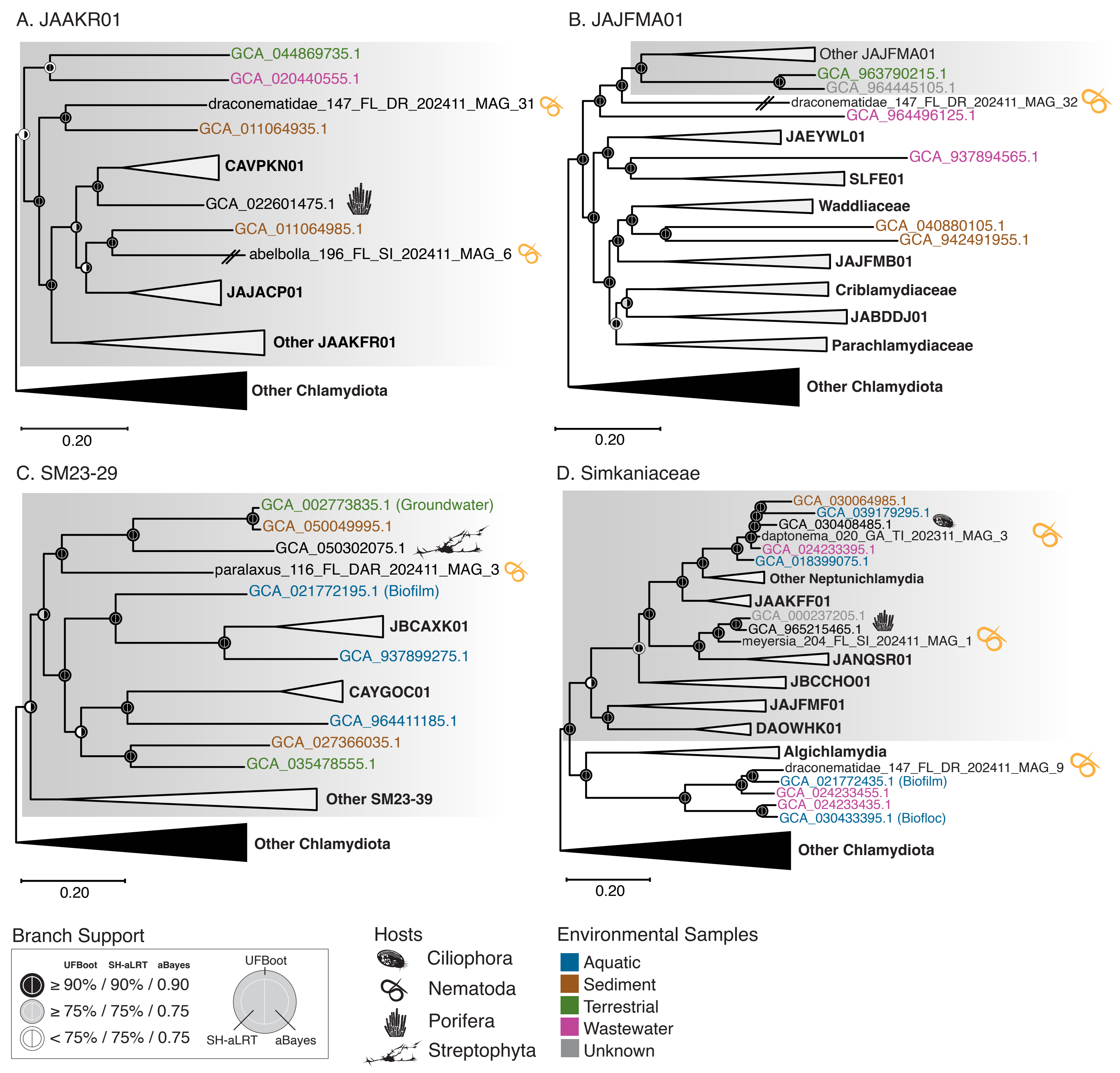

### Figure S5

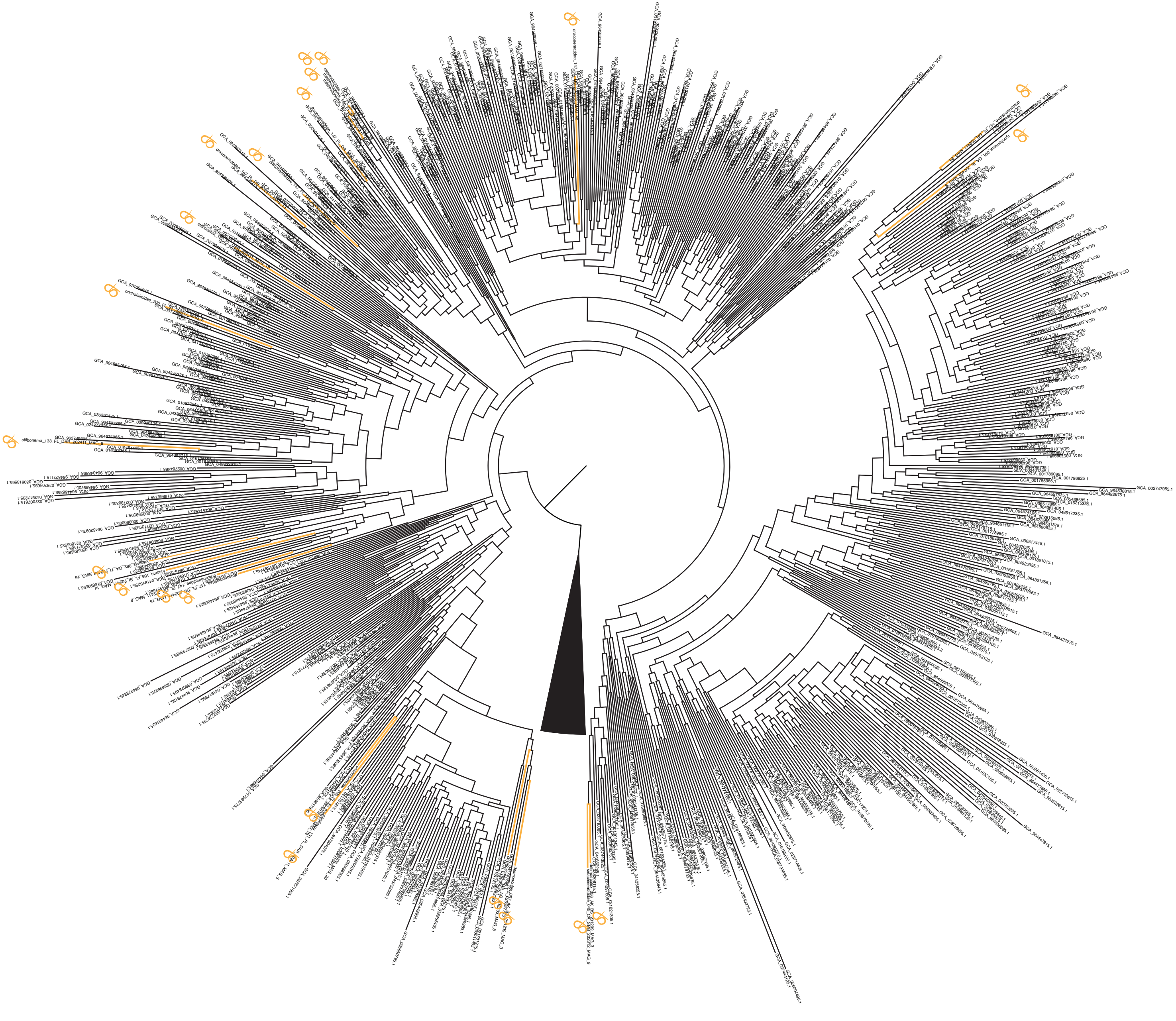

### Figure S6

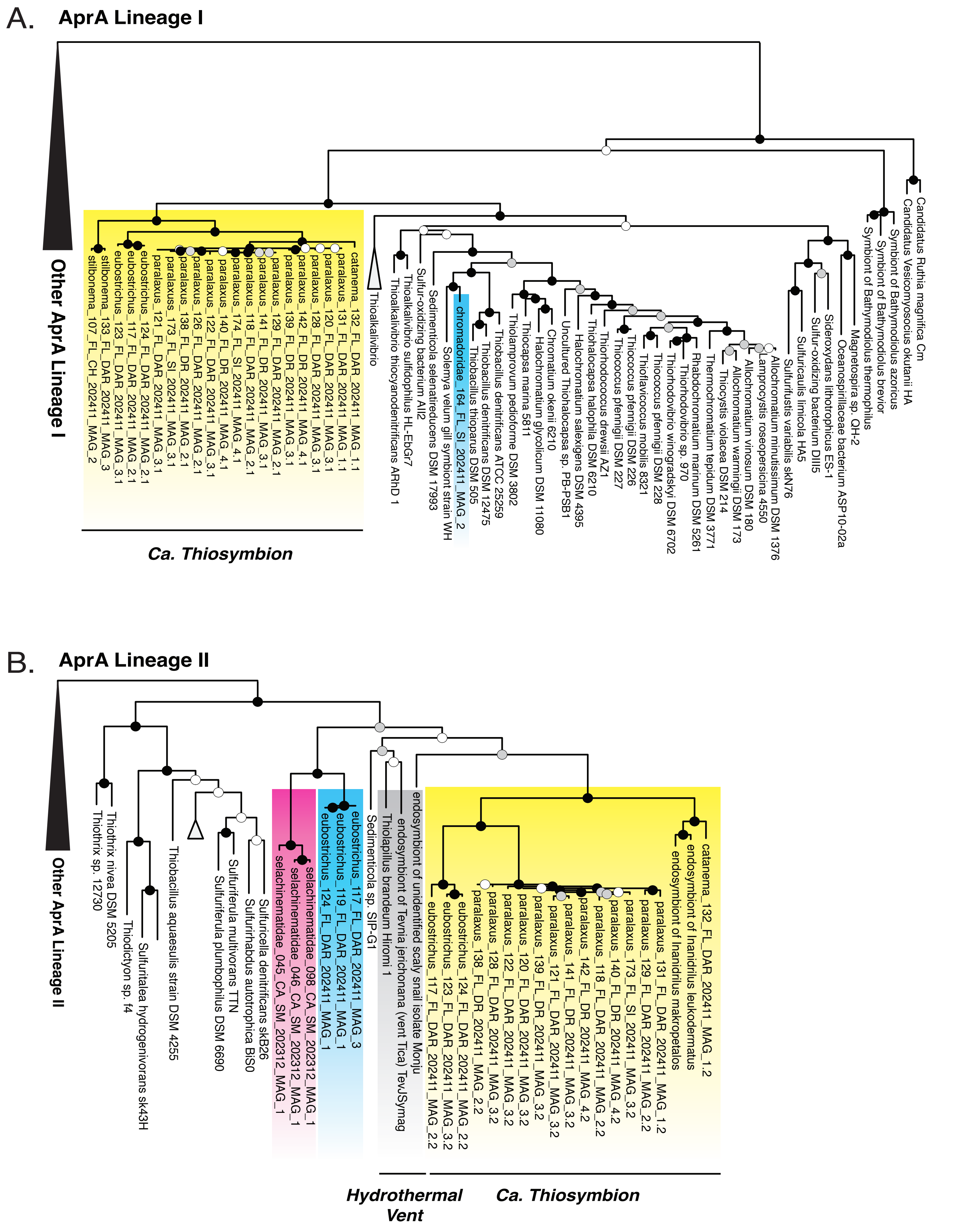

### Figure S7

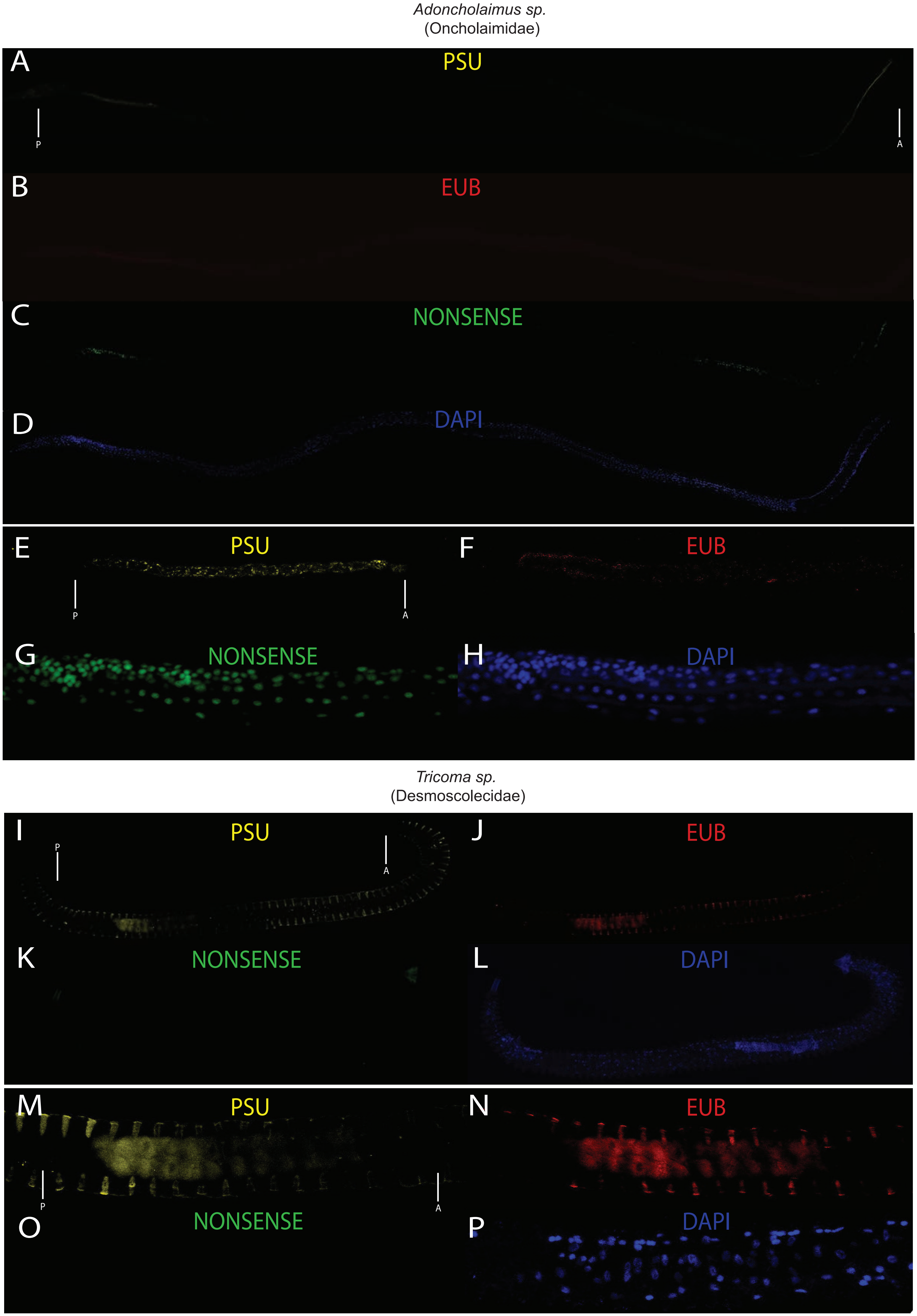
